# LBR and LAP2 mediate heterochromatin tethering to the nuclear periphery to preserve genome homeostasis

**DOI:** 10.1101/2024.12.23.628302

**Authors:** Renard Lewis, Virginia Sinigiani, Krisztian Koos, Cristiana Bersaglieri, Caroline Ashiono, Raffaella Santoro, Constance Ciaudo, Peter Horvath, Puneet Sharma, Ulrike Kutay

## Abstract

In most eukaryotic cells, euchromatin is localized in the nuclear interior, whereas heterochromatin is enriched at the nuclear envelope (NE). This conventional chromatin organization is established by heterochromatin tethering to the NE, however its importance for cellular homeostasis is largely unexplored. Peripheral heterochromatin localization relies on redundant NE-tethering systems. One tether is constituted by the lamin B receptor (LBR) in mammals, but the enigmatic nature of the other tethers has hampered functional analyses. Here we demonstrate that the downregulation of abundant, ubiquitous NE proteins can induce the global detachment of heterochromatin from the NE. Among these factors, we identify LBR and LAP2 as major players in bulk heterochromatin attachment to the NE in pluripotent and differentiated mammalian cells. Their loss leads to repositioning of heterochromatin to the nuclear interior, changes in chromatin accessibility, deregulation of gene expression including activation of antiviral innate immunity, and defects in cell fate determination.

---

Heterochromatin is positioned at the nuclear periphery and around nucleoli, whereas euchromatin is in the nuclear interior and in vicinity of nuclear pore complexes^1–7^. This ‘conventional’ nuclear architecture is an almost universal feature of eukaryotic nuclei and achieved by tethering of heterochromatin to the nuclear envelope (NE)^8,9^.

In metazoan cells, peripheral heterochromatin resides in proximity to the nuclear lamina, a protein meshwork formed by membrane proteins of the inner nuclear membrane (INM) and nuclear lamins, the archetype intermediate filament proteins^10–12^. The major lamin isoforms of mammalian cells are two A-type lamins, lamins A and C, encoded by the same gene, and two B-type lamins, lamins B1 and B2, encoded by separate genes. The genomic regions in proximity to the nuclear lamina are referred to as lamina-associated domains (LADs) and have been mapped by genome-wide DNA modification approaches^13–17^. LADs are between 0.1 to 10 Mb in size, gene-poor, AT-rich, enriched in epigenetic heterochromatic marks, replicate late in S phase, and correspond by-and-large to the B compartments that were defined by Hi-C^16,18,19^. Integrative LAD mapping has assigned up to 40% of genomic regions to LADs, but many LADs dynamically reshuffle into nucleolus-associated domains (NADs) upon cell division^5,16,19–23^.

An outstanding question in the field concerns the mechanisms that govern the peripheral positioning of mammalian heterochromatin. The prevailing hypothesis posits that heterochromatin attachment to the NE of mammals is accomplished by the collaborative effort of two molecular assemblies, referred to as the ‘A and B-type tethers’^8^. The A-type tether is organized by A-type lamins and functionalized by unknown proteins of the INM, whereas the B-type tether is constituted by the INM protein lamin B receptor (LBR)^8^ that binds lamin B1^24^. The presence of either the A-type or B-type tether is sufficient to maintain heterochromatin at the periphery, with the dominance of the respective tethers differing according to the differentiation stage of cells^8^.

Peripheral positioning of genes in heterochromatin has been implicated in the regulation of gene expression and thereby cell fate specification during differentiation and development^7,12,16,17,25–27^. Artificial tethering of genomic loci to the nuclear lamina can lead to their repression^28,29^, indicating that peripheral localization can play an active role in gene silencing. Systematic studies in mammalian cells support the view that LADs impede transcriptional activity and store silent genomic loci in the absence of their activating factors^30,31^, although certain peripherally tethered loci can escape repression^15,32,33^. Conversely, disruption of heterochromatin, local chromatin decondensation or transcriptional activation of genes can cause the relocation of genes into the nuclear interior^34–36^, suggesting that chromatin structure influences the peripheral localization of genomic loci.

Given all the evidence, anchoring heterochromatin at the nuclear periphery seems to be a fundamental feature of metazoan genome organisation. However, a full understanding of the mechanisms governing heterochromatin localization to the INM as well as of the relationship between its sequestration at the NE and its functions are currently lacking, likely because of the innate redundancy of the tethering mechanism. Besides its potential role in repressing genes found in LADs, it is not clear if other aspects of genome homeostasis and nuclear function are jeopardized when heterochromatin is detached from the nuclear periphery. We therefore set out to identify which mammalian INM proteins contribute to bulk heterochromatin tethering to the NE and study the consequences of changing heterochromatin distribution. We show that the downregulation of abundant, ubiquitously expressed NE proteins leads to the detachment of heterochromatin from the nuclear periphery in a range of differentiated mammalian cell lines and embryonic stem cells (ESCs). Among these factors, we identify major contributions from LBR and LAP2 in tethering heterochromatin to the NE in differentiated and pluripotent mammalian cells. Loss of these tethers causes changes in gene expression, chromatin organization, histone methylation, transposon silencing, splicing, and cell differentiation, highlighting the functional importance of peripheral heterochromatin localisation.

## Results

### Depletion of abundant NE proteins causes detachment of constitutive heterochromatin

The attachment of constitutive heterochromatin to the NE in mammalian cells is assumed to be redundantly governed by LBR and one or several undefined INM proteins^8^. To identify NE–heterochromatin tethers, and considering their potential redundancy, we employed a reverse genetics approach and collectively depleted 12 abundant, ubiquitously expressed NE factors using a mixture of siPOOLs (Fig. 1a). Each individual siPOOL is composed of about 30 siRNAs that target a single mRNA of interest and are effective at a very low concentration, minimizing off-target effects^37^. The targeted factors comprised both A-type lamins (lamin A and lamin C), the B-type lamins (lamin B1 and lamin B2) and eight INM proteins. The chosen INM proteins (Fig. 1a) included LBR, the four human LEM (<u>L</u>AP2β-<u>E</u>merin-<u>M</u>AN1) domain proteins Emerin (EMD), LAP2, LEMD2, and MAN1, since LEM domain factors have been implicated in heterochromatin positioning in diverse eukaryotes^38–44^, the SUN domain proteins SUN1 and SUN2 known to be involved in diverse aspects of chromatin organization^45^, and the chromatin-binding INM protein LAP1^46^. Note that only 11 si-POOLs were needed (si-11) for depletion of the 12 factors, since lamin A and lamin C mRNAs are largely identical and can be depleted by the same siPOOL.

**Fig. 1.**
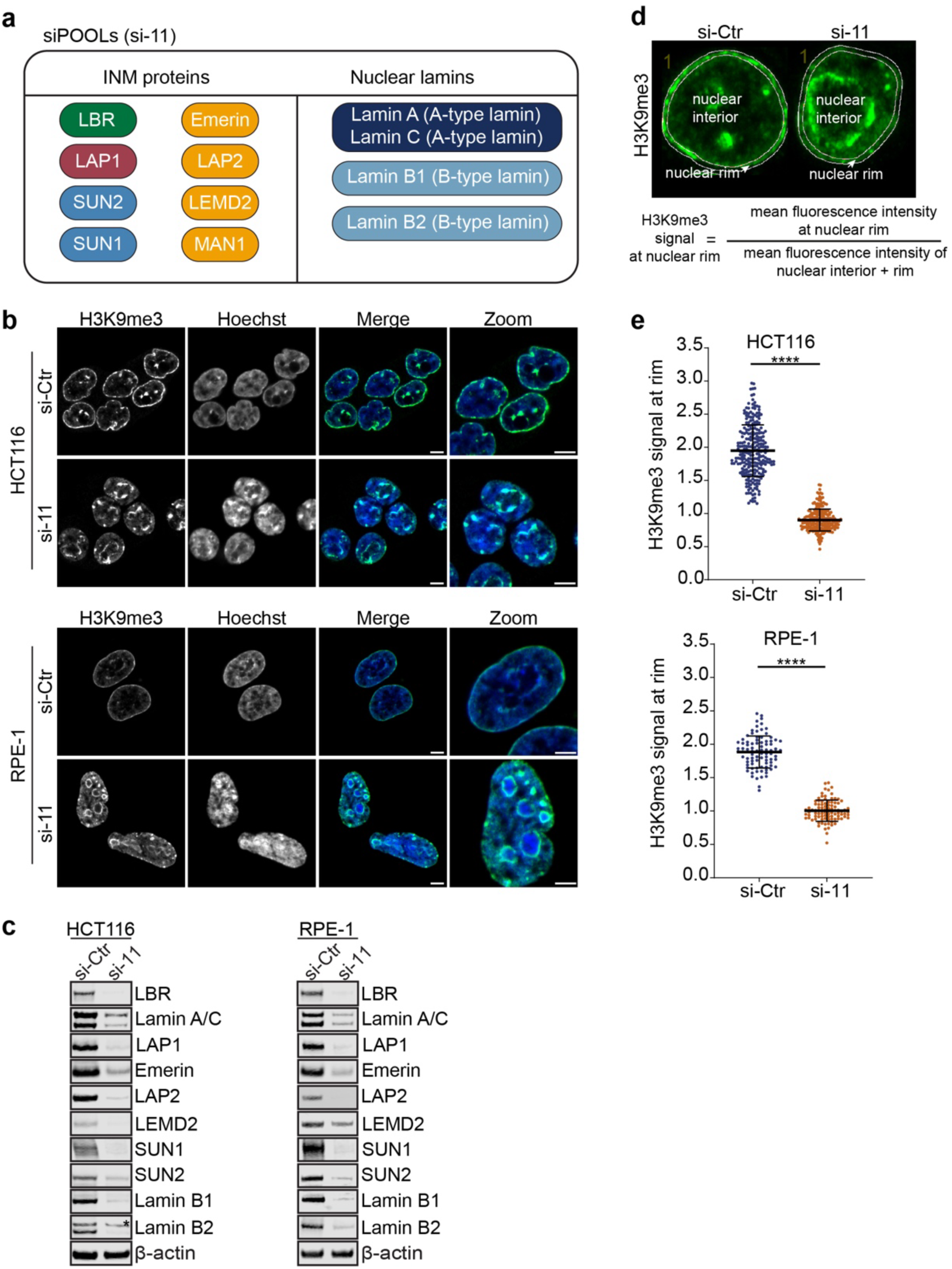
H3K9me3-stained heterochromatin is detached from the nuclear periphery of human cells upon depletion of 8 INM proteins and the nuclear lamins. **a**, Selected set of 12 NE factors (LBR, SUN1, SUN2, LAP1, Emerin, LAP2, LEMD2, MAN1, lamin A, lamin C, lamin B1 and lamin B2) targeted by a combination of siPOOLs (si-11) **b**, Representative confocal images of HCT116 and RPE-1 cells either treated with control siRNA (si-Ctr) or a combination of 11 siPOOLs for 72 h, fixed and subjected to immunofluorescence using an anti-H3K9me3 antibody. DNA was stained with Hoechst. Scale bars, 5 μm **c**, Immunoblots confirming the downregulation of select NE proteins upon si-11 depletion. Asterisks shows an unspecific band in the HCT116 lamin B1 blot. **d**, Quantification procedure of the fraction of H3K9me3-positive heterochromatin at the nuclear rim. H3K9me3 intensity was quantified in a ∼ 0.4 μm-wide ring at the nuclear periphery and expressed as the ratio of the average integrated fluorescence intensities at the nuclear rim over the entire nuclear area (nuclear rim + nuclear interior). **e**, Quantification of the H3K9me3 signal at the nuclear rim in HCT116 cells (n ≥ 227) and RPE-1 cells (n ≥ 86) treated as in (b) (*p ≤ 0.05, **p ≤ 0.01, ***p ≤ 0.001, ****p≤ 0.0001, Welch’s t-test, N = 3). Error bars, SD.

To examine how depletion of the chosen factors affects heterochromatin positioning, we initially analyzed HCT116 cells, a differentiated near-diploid human colorectal carcinoma cell line (Fig. 1b). Heterochromatin was visualized by immunofluorescence using an antibody against the epigenetic histone modification H3K9me3 that marks constitutive heterochromatin and is enriched in LADs^16,47,48^. Immunoblotting of a large subset of the chosen factors confirmed their efficient downregulation (Fig. 1c), which was further confirmed by RNA-seq (Extended Data Fig. 1a, Extended Data Table 1). HCT116 cells treated with control siRNA displayed the typical peripheral enrichment of H3K9me3 and a few intranuclear heterochromatin foci (Fig. 1b). In contrast, cells treated with the si-11 pool showed a strong decrease in peripheral H3K9me3 staining and a striking accumulation of intranuclear heterochromatin clusters (Fig. 1b). Comparison with the Hoechst signal indicated that H3K9me3-positive foci were in the vicinity of nucleoli, indicating that loss of peripheral tethering might convert some LADs into NADs (Fig.1b).

To quantitatively assess the rearrangement of peripheral heterochromatin, we delineated the nuclear contours based on the Hoechst signal and measured the mean integrated density of H3K9me3 in a ∼ 0.4 μm wide ring at the nuclear periphery relative to the entire the nucleus (Fig. 1d). These quantifications showed a significant decrease in the H3K9me3 signal at the NE upon depletion of the 12 targeted NE proteins in both HCT116 and RPE-1 cells (Fig. 1e). Analysis of H4K20me3 and H3K27me3 that decorate constitutive and facultative heterochromatin, respectively, further supported the global loss of heterochromatin from the NE (Extended Data Fig. 1b,c). The repositioning of H3K9me3-positive heterochromatin into the nuclear interior was also observed in human RPE-1 cells, a diploid retinal pigment epithelial cell line (Fig. 1b).

We also confirmed these findings in mouse cells, examining both differentiated NIH/3T3 fibroblasts and mouse embryonic stem cells (mESCs) (Extended Data Fig. 1d,e), indicating that the involved tethers are conserved between humans and rodents. Taken together, depletion of nuclear lamins and eight abundant INM proteins is sufficient to induce the detachment of heterochromatin from the NE in a variety of mammalian cell lines.

### Depletion of LBR and lamin A/C is insufficient to globally release constitutive heterochromatin from the NE

Landmark studies in the rod cells of mice have demonstrated that the absence of LBR and lamin A/C is sufficient to cause nuclear chromatin inversion in this particular cell type^8^. Therefore, we next set out to examine whether the downregulation of LBR and lamin A/C was potentially sufficient for heterochromatin detachment from the NE in HCT116 cells. Since siRNA-mediated downregulation of A-type lamins was less efficient than that of LBR (Fig. 1c), we first generated *LMNA* KOs HCT116 cells (Extended Data Fig. 2a). Then, using both HCT116 wild-type (WT) and *LMNA* KO cell lines, we compared the effect of LBR depletion with that of all selected NE proteins on peripheral heterochromatin positioning using an adapted pool of siRNAs (si-10; excluding the siPOOL for lamin A/C) (Extended Data Fig. 2b–d). The individual loss of either LBR (siP-LBR) or lamin A/C (in the *LMNA* KO cells) had discernible but minor effects on peripheral heterochromatin positioning, evident by the slightly reduced enrichment of H3K9me3-positive chromatin at the NE and the formation of small intranuclear heterochromatin foci. The combined loss of lamin A/C and LBR clearly exacerbated these defects. However, only the combined depletion of all other 10 NE factors in HCT116 *LMNA* KO led to a gross reduction in the level of peripheral heterochromatin and the formation of large intranuclear chromatin clusters (HCT116 LMNA KO, si-10). This indicates that the absence of LBR and lamin A/C is insufficient to release heterochromatin from the NE to the same extent as the loss of all 12 proteins, indicating that there are factors other than LBR and lamin A/C contributing to bulk heterochromatin tethering in HCT116 cells. Intriguingly, even in the presence of lamin A/C (HCT116 WT), the loss of the other 10 proteins still had a stronger effect on peripheral heterochromatin accumulation than loss of LBR alone, further suggesting a contribution of other factors (Extended Data Fig. 2b-d).

### Identification of mammalian INM proteins that anchor heterochromatin

To identify which other factors promote heterochromatin tethering to the NE, we set out to deconvolute the complexity of the siPOOL depletion experiments in the HCT116 *LMNA* KO background. We took a systematic approach to define the loss of which protein(s) elicit(s) an effect on peripheral heterochromatin positioning (Fig. 2a), initially breaking apart the si-10 pool into five bins (LBR, B-type lamins, LEM domain proteins, SUN protein, LAP1), grouping factors belonging to the same protein families. As expected, we observed a significant but incomplete release of peripheral H3K9me3-marked heterochromatin from the NE upon depletion of LBR (Step 1, Fig. 2a, Extended Data Fig. 3a,b,g). Next, we downregulated each of the four remaining groups on top of LBR, which revealed a contribution of the LEM domain protein family (Step 2, Extended Data Fig. 3c,d,g). To pinpoint which LEM domain family members contribute, we next compared co-downregulation of LBR with each of the individual LEM domain proteins (LAP2, Emerin, LEMD2, or MAN1) with the LEM family pool (Step 3, Fig. 2b,c,g). We observed a significant effect of LAP2, the depletion of which exacerbated the effect of LBR downregulation. Cells co-depleted of LBR and LAP2 contained large intranuclear heterochromatin clusters and by-and-large lacked peripheral H3K9me3-positive heterochromatin at the nuclear rim. Depletion of none of the other LEM domain proteins caused any significant changes. Finally, we tested the remaining three groups (B-type lamins, SUN proteins, LAP1) in conjunction with combined LBR and LAP2 downregulation (Step 4, Extended Data Fig.3 e–g). This did not reveal any further enhancement of the phenotype induced by co-depletion of LBR and LAP2.

**Figure 2.**
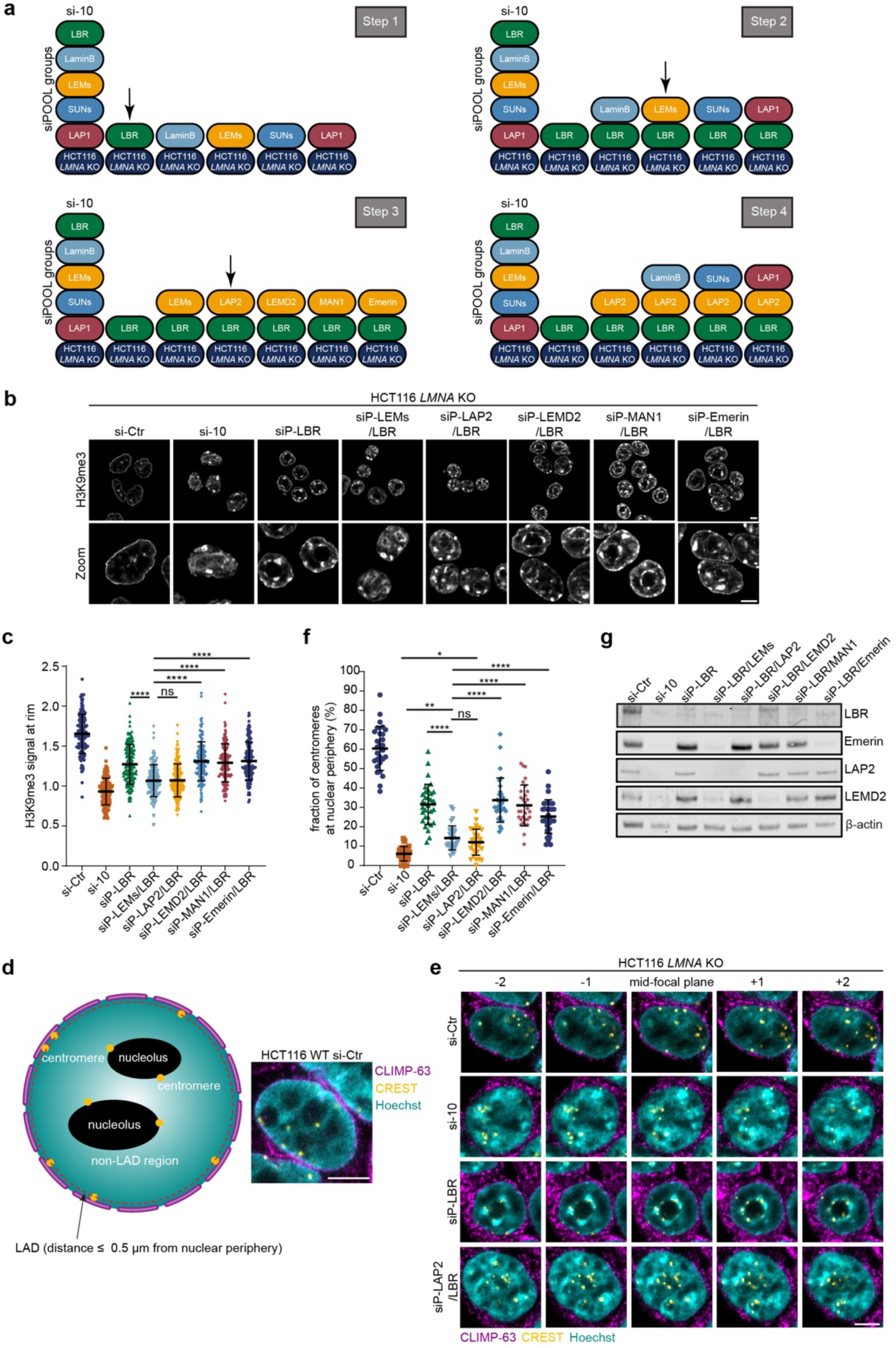
The INM proteins LBR and LAP2 promote the peripheral positioning of H3K9me3-positive heterochromatin. **a**, Schematic depiction of the deconvolution approach performed in HCT116 *LMNA* KO cells. The proteins targeted by si-10 were initially distributed into five groups: LBR, B-type lamins (lamin B1, lamin B2), LEM family (LAP2, Emerin, LEMD2, MAN1), SUN proteins (SUN2, SUN2) and LAP1. The corresponding sets of siPOOLs were first individually tested (Step 1), identifying LBR. In the following steps, the contributions of additional factors were evaluated as outlined. Arrows indicate the major additional factors affecting peripheral heterochromatin identified in each step **b**, Representative confocal images of H3K9me3-positive heterochromatin in HCT116 *LMNA* KO cells treated with si-Ctr or the indicated combination of siPOOLs for 72 h (Step 3 of deconvolution), identifying LAP2. Scale bars, 5 μm. **c**, Quantification of the H3K9me3 signal at the nuclear rim for (b) (n ≥ 133, *p ≤ 0.05, **p ≤ 0.01, ***p ≤ 0.001, ****p≤ 0.0001, one-way ANOVA, N = 3). Error bars, SD. **d**, Quantification of centromeres at the NE. In wild-type cells, a fraction of centromeres is localized at the nuclear periphery, whereas internal centromeres are enriched around nucleoli. To quantify the fraction of centromeres in the vicinity of the NE, fixed cells were co-stained for centromeres (CREST antibody), CLIMP-63 (ER/NE network) and DNA (Hoechst). Z-stacks were acquired by confocal microscopy throughout the cell volume. All centromeres at a distance of ≤ 0.5 μm from the nuclear periphery were considered to be positioned in LADs, which extend up to 1 μm into the nuclear interior^20^. The fraction of centromeres at the NE was determined as the ratio between the number of peripheral centromeres and all identified centromeres. Scale Bar, 5 μm **e**, Representative confocal images of CREST, CLIMP-63 and Hoechst (DNA) throughout five microscopy slices in HCT116 *LMNA* KO cells treated with si-Ctr or the indicated combination of siPOOLs for 72 h (Step 3 of deconvolution), identifying LAP2. Scale bars, 5 μm. **f**, Quantification of the proportion of centromeres at the nuclear periphery in cells treated with si-Ctr or the indicated combinations of siPOOLs for 72 h (n ≥ 28, *p ≤ 0.05, **p ≤ 0.01, ***p ≤ 0.001, ****p≤ 0.0001, one-way ANOVA, N= 3). Error bars, SD. **g,** Immunoblotting of LBR, LAP2 and select other NE proteins to determine the efficiency of siRNA-induced protein depletion.

As a complementary readout for the position of constitutive heterochromatin, we analysed centromeres, which are embedded in constitutive heterochromatin^49^. A fraction of centromeres is known to be positioned at the nuclear periphery throughout the cell cycle, whereas centromeres located in the nuclear interior are adjacent to nucleoli^50–52^. We observed that about 60% of centromeres were localized at the nuclear periphery in siCtr-treated HCT116 WT cells (Extended Data Fig. 4a), in agreement with recent analyses in HeLa cells^52^. Interestingly, the absence of lamin A/C in the HCT116 *LMNA* KO cells did not perturb peripheral centromere positioning compared to WT HCT116 cells, whereas RNAi-mediated depletion of the set of selected NE proteins by si-10 strongly diminished peripheral centromere localization, with fewer than 10% of centromeres remaining at the nuclear periphery (Extended Data Fig. 4a).

Repetition of the stepwise deconvolution process (Fig. 2a) using centromere positioning as a readout confirmed and extended the conclusions drawn from H3K9me3 staining (Fig. 2d–f, Extended Data Fig. 4b–d). Step 1 revealed that depletion of LBR, LEM-domain proteins and LAP1 reduced the proportion of centromeres at the nuclear periphery, with values dropping from about 60% (si-Ctr) to 25%, 35% and 50%, respectively (Extended Data Fig. 4b). Co-depletions of LBR with the other groups of NE factors in step 2 of the deconvolution process demonstrated that the group of LEM-domain proteins had the strongest effect on the peripheral positioning of centromeres when co-depleted with LBR, followed by LAP1 and some minor effect of the B-type lamins (Extended Data Fig. 4c). Strikingly, when breaking apart the group of LEM domain proteins (Step 3), co-depletion of LBR and LAP2, but not of other LEM-domain proteins, strongly affected peripheral centromere positioning, reducing the fraction of centromeres at the nuclear periphery to about 10%, akin to co-depletion of LBR and all LEMs (Fig. 2f). This suggests that LAP2 and LBR are chiefly responsible for peripheral centromere positioning, mirroring what was observed with the H3K9me3 readout. Notably though, downregulation of LBR and LAP2 was not fully sufficient to recapitulate the si-10 phenotype (Fig. 2f), suggesting that additional factors might support centromere tethering. Further co-depletions of LBR, LAP2 and the remaining groups of NE factors (step 4) revealed that loss of LAP1 causes a small but significant effect on centromere anchoring (Extended Data Fig. 4d). Collectively, these experiments identify the INM proteins LBR and LAP2 as the major players in tethering of constitutive heterochromatin to the NE in *LMNA* KO cells, with a minor effect of LAP1 on centromeres that remains to be further explored.

### The membrane-anchored isoforms of LAP2 promote heterochromatin attachment to the NE

Human cells express four isoforms of LAP2 (LAP2α, β, γ and δ, Fig. 3a), of which LAP2α and LAP2β are the most abundant^53^. All isoforms can interact with chromatin via their N-terminal LEM-like and LEM-domains. LAP2α lacks a transmembrane domain and localizes to the nucleoplasm, where it promotes binding of lamin A/C to heterochromatin^54^, whereas the other isoforms are embedded in the INM (Fig. 3a). Therefore, we asked whether it is indeed the membrane-bound isoforms that synergize with LBR in promoting heterochromatin localization to the NE, especially in cells that express the A-type lamins. To ensure the equal absence of LBR in all cells, we generated HCT116 *LBR* KO cells (Extended Data Fig. 5a). H3K9me3 localization in *LBR* KO cells was slightly perturbed, evident by the more prominent intranuclear heterochromatin foci and an uneven H3K9me3 distribution at the NE (Fig. 3b, si-Ctr). Additional downregulation of the other NE proteins including LAP2 caused a global detachment of heterochromatin from the NE, as expected (Fig. 3b,c, si-10b). The individual depletions of either LAP2 or lamin A/C in the HCT116 *LBR* KO cells caused similar effects on peripheral heterochromatin localization, accentuating the phenotype seen in the HCT116 *LBR* KO background.

**Figure 3.**
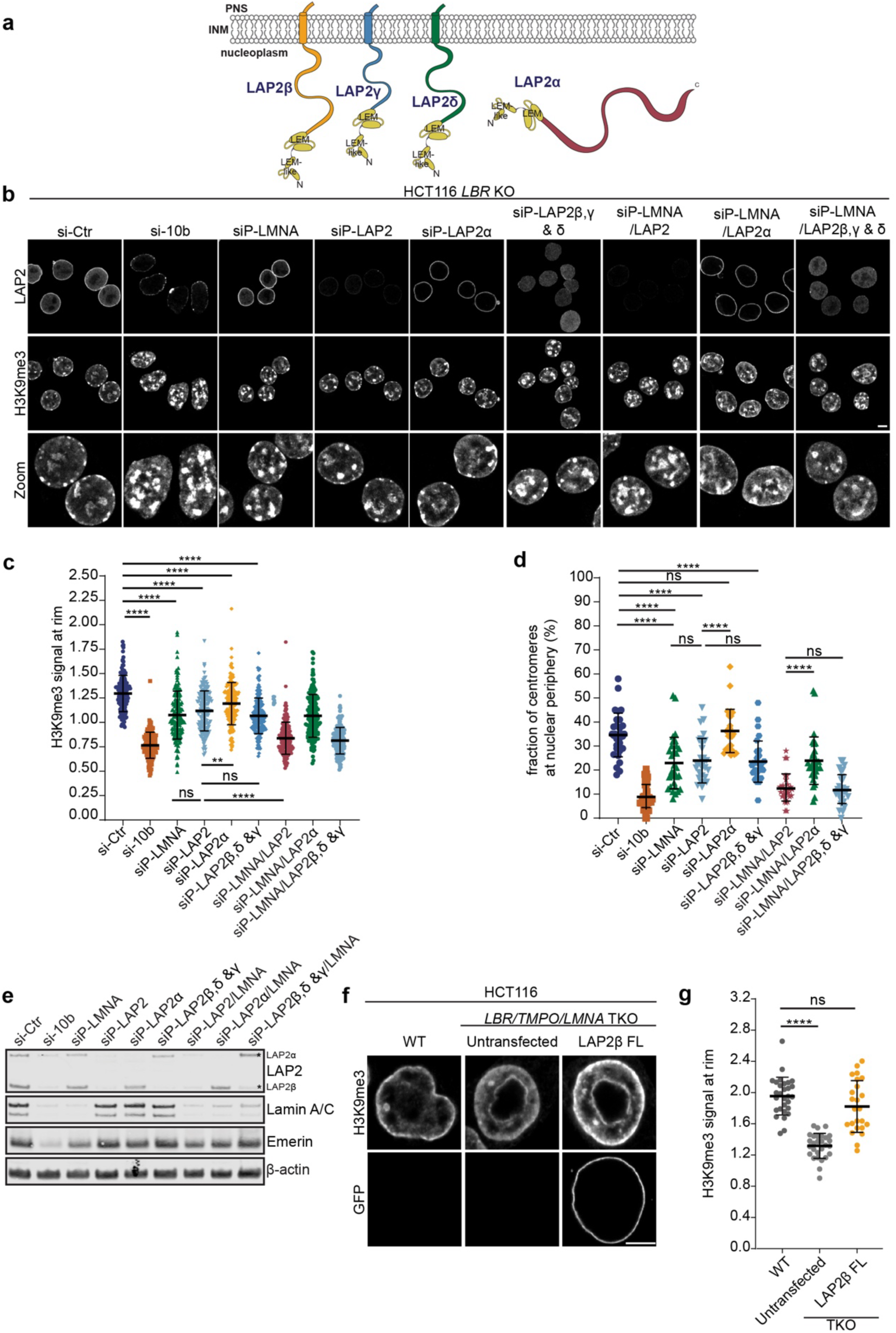
The INM-bound isoforms of LAP2 contribute to peripheral heterochromatin anchorage. **a**, Schematic depiction of the four isoforms of LAP2 expressed in human cells. Note that the presence of LAP2δ at the protein level is still contested. **b**, Representative confocal images of H3K9me3 localisation in HCT116 *LBR* KO cells treated with si-Ctr or the indicated combinations of siPOOLs for 72 h. si-10b refers: siPOOL mixture including siP-lamin A/C but not siP-LBR. Scale bars, 5 μm. **c**, Quantification of the H3K9me3 signal at the nuclear rim in (b) (n ≥ 137, *p ≤ 0.05, **p ≤ 0.01, ***p ≤ 0.001, ****p≤ 0.0001, one-way ANOVA, N = 3). Error bars, SD. **d**, Quantification of the proportion of centromeres at the nuclear periphery in cells treated as in (b) (n ≥ 27, *p ≤ 0.05, **p ≤ 0.01, ***p ≤ 0.001, ****p≤ 0.0001, one-way ANOVA, N = 3). Error bars, SD. **e**, Immunoblotting of LAP2 (detecting LAP2α and LAP2β, asterisks), lamin A/C and emerin to determine the efficiency of siRNA-induced protein depletion. **f**, Representative confocal images of H3K9me3 localisation in HCT116 WT and *LBR/TMPO/LMNA* TKO cells as well as of TKO cells expressing GFP-tagged LAP2β after 48 h of transient transfection. Scale bars, 5 μm. **g**, Quantification of the H3K9me3 signal at the nuclear rim in in (f) (n ≥ 24, *p ≤ 0.05, **p ≤ 0.01, ***p ≤ 0.001, ****p≤ 0.0001, one-way ANOVA, N = 3). Error bars, SD.

For isoform group-specific depletion of LAP2, we exploited siPOOLs designed to target the mRNAs of either LAP2α or all membrane-bound isoforms (LAP2β, γ and δ). These siRNAs pools displayed the expected specificity as evident by immunofluorescence and immunoblotting analyses (Fig. 3b, e). The depletion of the membrane-bound LAP2 isoforms mirrored the effect of downregulation of all LAP2 isoforms (siP-LAP2). It is noteworthy that the co-depletion of lamin A/C together with either all LAP2 or the membrane-bound LAP2 isoforms affected H3K9me3 localisation more strongly, indicating an A-type lamin-dependent contribution to heterochromatin tethering that remains to be defined. Analysis of centromere positioning confirmed that the loss of the membrane-bound LAP2 isoforms mimics the effect of total LAP2 depletion, whereas the loss of LAP2α does not (Fig. 3d).

We conclude that the INM-bound isoforms of LAP2 promote heterochromatin tethering to the NE. The role of LAP2 in heterochromatin attachment to the nuclear periphery was further confirmed by a rescue experiment exploiting a genome-edited HCT116 cell line in which we had ablated the expression of LBR, LAP2 and LMNA/C (*LBR/TMPO/LMNA* triple KO (TKO); see below and Extended Data Fig. 5f,g) and re-expressed LAP2β after transient transfection, which induced the enrichment of H3K9me3-positive chromatin at the NE (Fig. 3f,g).

### Loss of LBR and LAP2 affects chromatin organization and gene expression

Most differentiated eukaryotic cells position their heterochromatin at the nuclear periphery, however, the biological consequences of a global loss of peripheral heterochromatin have not been explored. Having a system in hand that allows inducing detachment of heterochromatin from the NE at will, we set out to determine the consequences of the loss of peripheral heterochromatin positioning on gene expression and genome organization.

Short-term depletion of NE proteins in WT HCT116 cells by RNAi (si-11, 72 h) affected gene expression very mildly, as revealed by RNA-seq (Extended Data Fig. 1a). Only a small number of genes became significantly deregulated, with 13 and 28 genes being down- and upregulated more than twofold, respectively. The 13 downregulated genes included 10 of the genes targeted by the si-11 depletion. To assign the observed changes to LAD or inter-LAD regions, we used an annotated lamin B1 DamID dataset for HCT116 cells^23^. Affected genes originally positioned in LADs were upregulated (Extended Data Fig. 6a). ATAC-seq, which measures the accessibility of genomic regions to a hyperactive Tn5 transposase^55^, demonstrated that only a small fraction of the genome (∼ 1.6%, LFC ≥ l1l) exhibited altered accessibility upon short-term depletion of NE tethers. Here, genomic regions known to be situated in LADs^23^ became largely inaccessible (Extended Data Fig. 6b,c; Extended Data Table 2), likely due to the collapse of peripheral chromatin into intranuclear heterochromatin clusters. Overall, we conclude that short-term effects of RNAi-induced heterochromatin release from the NE on gene expression are minor.

To examine the effects of prolonged detachment of heterochromatin from the NE, we generated KO cell lines that carry double deletions of the genes for LBR and LAP2 encoded by the *TMPO* gene (*LBR/TMPO* double KO (DKO)) or the combined deletion of the genes encoding for LBR, LAP2 and LMNA/C (*LBR/TMPO/LMNA* triple KO (TKO)) (Extended Data Figure 5b-g). We first analysed the localisation of heterochromatin in these cell lines using antibodies for H3K9me3 and H3K27me3 that decorate constitutive and facultative heterochromatin, respectively (Fig. 4a,b). Consistent with the RNAi experiments, we observed a decrease in the staining of H3K9me3 at the nuclear periphery in both genetic backgrounds, accompanied by the appearance of large clusters of collapsed intranuclear heterochromatin. Similarly, H3K27me3-positive heterochromatin was detached from the NE, indicating that H3K27me3-marked facultative heterochromatin is also affected.

**Figure 4.**
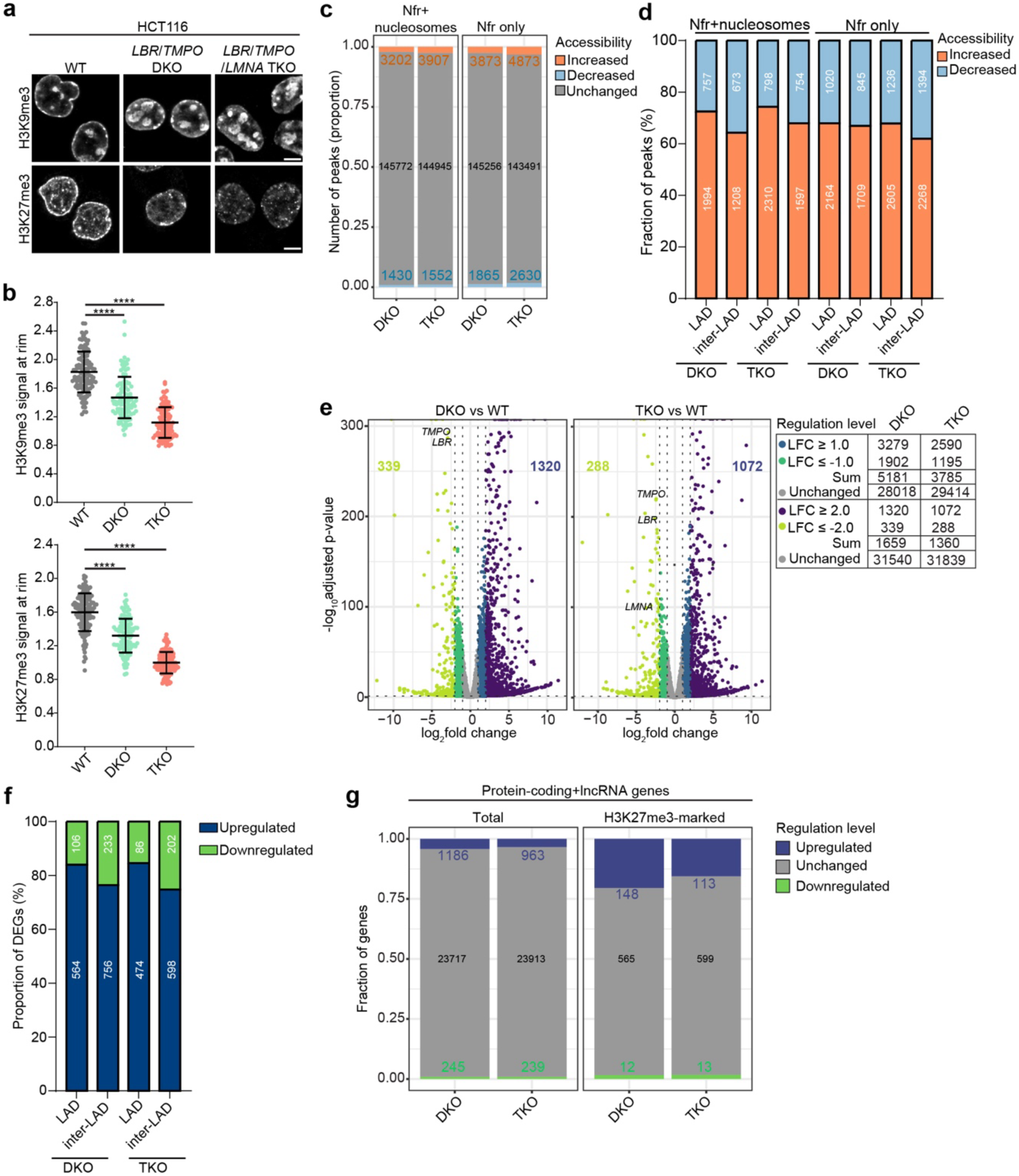
Genomic deletion of LBR and LAP2 severely perturbs chromatin organization and gene expression in HCT116 cells. **a**, Representative confocal images of H3K9me3- and H3K27me3-positive heterochromatin in HCT116 *LBR/TMPO* DKO and *LBR/TMPO/LMNA* TKO. Scale bars, 5 μm. **b**, Quantification of the H3K9me3 signal (n ≥ 101, *p ≤ 0.05, **p ≤ 0.01, **p ≤ 0.001, ****p≤ 0.0001, one-way ANOVA, N = 3) and H3K27me3 signal (n ≥ 111, *p ≤ 0.05, **p ≤ 0.01, **p ≤ 0.001, ****p≤ 0.0001, one-way ANOVA, N = 3) at the nuclear rim in (a). Error bars, SD. **c**, Proportion of ATAC-seq peaks in nucleosome-free regions (Nfr) or in both Nfr and nucleosomes-containing regions (Nfr+nucleosomes) with increased or decreased accessibility in *LBR/TMPO* DKO and *LBR/TMPO/LMNA* TKO cells (N = 3). All loci with a log_2_-fold change ≥ l2l and an adjusted p-value ≤ 0.05 were considered significant. **d**, Bar graphs highlighting the proportion of Nfr+nucleosomes and Nfr only ATAC-seq peaks with increased or decreased accessibility in regions designated as LADs in *LBR/TMPO* DKO and *LBR/TMPO/LMNA* TKO cells. **e**, Volcano plots of differential gene expression identified by RNA-seq (HCT116 *LBR/TMPO* DKO and *LBR/TMPO/LMNA* TKO vs. HCT116 WT cells, N = 3). Number of genes with an adjusted p-value ≤ 0.05 are marked as up- or downregulated if log_2_-fold change ≥ l2l in the Volcano blots. Table summarizes two- and fourfold changes in gene expression. **f**, Bar graphs showing the proportions of significantly up- or downregulated genes (log_2_-fold change ≥ l2l) within genomic regions originally positioned in LADs or inter-LADs in *LBR/TMPO* DKO and *LBR/TMPO/LMNA* TKO cells. **g**, Bar graphs showing the fraction of deregulated protein-coding and lncRNA genes identified by RNA-seq (e, log_2_- fold change ≥ l2l) that are enriched for H3K27me3 marked regions (ENCSR810BDB) within 500 nt of their transcription start sites.

ATAC-seq revealed widespread changes in genome accessibility, with about 14% of the detected genomic regions changing accessibility in both the DKO and TKO cells more than twofold and about 3% more than fourfold (Fig. 4c; Extended Data Table 2). In the *LBR/TMPO/LMNA* TKO, these changes were slightly more pronounced, consistent with the even more complete release of heterochromatin from the nuclear periphery. Surprisingly, of the affected loci, about 60% to 75% displayed increased accessibility, both in regions originally positioned in LADs or inter-LADs (Fig. 4d), indicating that genes in LADs were not preferentially affected by the long-term changes in chromatin accessibility.

Bulk RNA-seq demonstrated that the deletion of heterochromatin anchors caused widespread changes to the transcriptome. Hundreds of genes were deregulated by more than fourfold (1’659 genes in total in the DKO (1320 up, 339 down) and 1’360 genes total in the TKO (1’072 up, 288 down) (Fig. 4e, Extended Data Fig. 6d, Extended Data Table 1). Nearly 5% of all detected protein-coding RNAs and lncRNAs were affected. Mirroring the changes in ATAC-seq, a larger fraction of genes showed an upregulation in gene expression, with about 80% of the changed transcripts being significantly increased in abundance in both cell lines, both in LAD and inter-LAD regions (Fig. 4f). Further analysis revealed a moderate but highly significant correlation between the RNA-seq and ATAC-seq data (Extended Data Fig. 6e; Extended Data Table 3)). Notably, when comparing the single KO, DKO, and TKO genotypes (Extended Data Fig. 6d), deletion of *LBR* alone had already major effects. The effects of *LBR* and *TMPO* deletion were synergistic when considering downregulated genes, consistent with the notion that the combined loss of LBR and LAP2 leads to a greater extent of heterochromatin collapse, potentially burying genes and augmenting gene expression.

To determine whether the upregulation in gene expression and increased chromatin accessibility involve defects in the maintenance of repressive heterochromatin marks upon release of chromatin from the nuclear periphery, we used quantitative immunoblotting. Strikingly, the levels of H3K27me3 were drastically reduced. We also observed a reduction in H3K9me2, whereas H3K9me3 and H4K20me3 remained nearly unaffected (Extended Data Fig. 6f,g). The reduction in H3K27me3 levels suggest that derepression of H3K27me3-marked facultative heterochromatin might contribute to the prominent upregulation of gene expression. Comparison with published H3K27me3 ChIP data revealed that about 20% and 16% of genes with H3K27me3 enriched in their promoters in HCT116 WT cells (ENCSR810BDB) were fourfold upregulated in the DKO and TKO, respectively, indicating that derepression of H3K27me3-marked facultative heterochromatin contributes to the massive upregulation of gene expression (Fig. 4g). Notably, H3K9me2 and H3K9me3 were affected to different levels, which might point at unique roles for both modifications, a hypothesis for which evidence is emerging in other model organisms^56,57^.

### The persistent loss of heterochromatin tethers leads to an upregulation of an antiviral immune response

The deregulation of gene expression in the KO cells might present a combination of different classes of changes: (1) genes that are directly deregulated by their repositioning, (2) genes that are indirectly affected by deregulation of class 1 genes, and (3) genes that are part of the cell’s concerted response to genome-wide deregulation. To discern functionally linked clusters of factors among the upregulated genes that might be part of the cellular response to the inflicted changes in chromatin organization and gene expression (class 3), we performed Gene Ontology (GO) term analysis (Fig. 5a, Extended data Fig. 7a). Interestingly, considering genes that are upregulated more than fourfold (Fig. 4e), GO terms pertaining to antiviral defence, chemokine signalling, and extracellular matrix organization were the most prominent (Fig. 5a).

**Figure 5.**
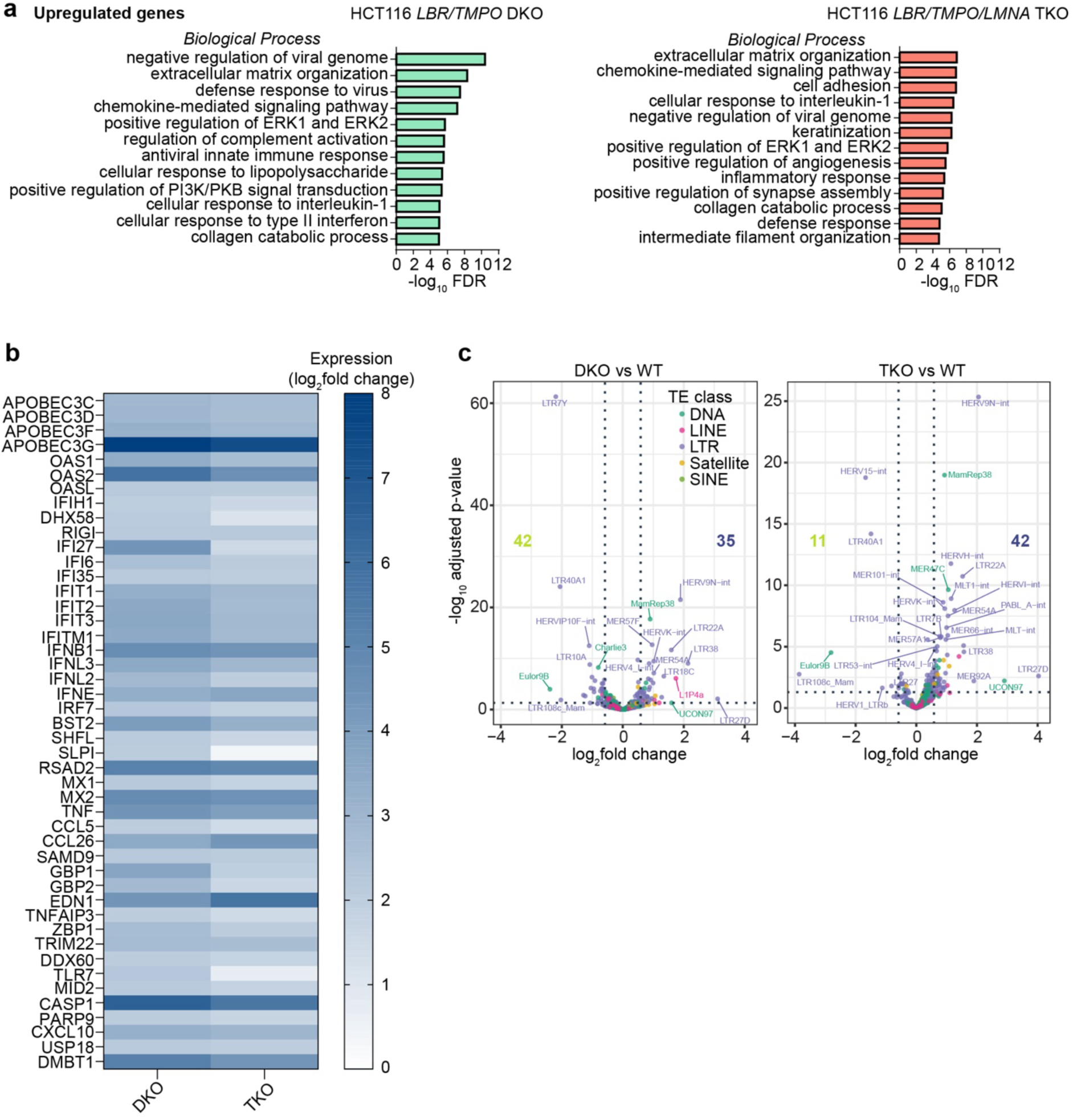
Genomic deletion of LBR and LAP2 induces clusters of genes linked to antiviral defence and innate immunity. **a**, Bar graph of Gene Ontology (GO) term analysis (‘Biological process’ category) of significantly upregulated genes with a log_2_-fold change ≥ 2 in HCT116 *LBR/TMPO* DKO and *LBR/TMPO/LMNA* TKO cells. Fischer’s exact test was used to determine overrepresented terms. **b,** Heatmap of innate immunity genes upregulated in the KO backgrounds and part of the identified GO terms ‘negative regulation of viral genome’ and ‘defense response to virus’. **c**, Volcano plots of RNA expressed from transposable elements identified by TETranscripts for *LBR/TMPO* DKO and *LBR/TMPO/LMNA* TKO cells vs HCT116 WT cells (N = 3, dashed lines indicate changes with p-value ≤ 0.05 and log_2_-fold change ≥ l0.585l). DNA transposon (DNA), Long interspersed nuclear elements (LINE), Long terminal repeat retrotransposon (LTR), Satellite repeats (Satellite), Small interspersed nuclear elements (SINE).

STRING analysis of the top 200 upregulated protein coding genes (Extended Data Fig. 7b) identified a cluster comprising prominent interferon-inducible innate immunity factors and cytokine genes including interferon beta 1 (IFN1B). Upregulated innate immunity factors (Fig. 5b) included the cytosolic dsRNA sensors^58^ RIG-1(DDX58, 4.5-fold in DKO) and MDA-5(IFIH1, 4.1-fold), oligoadenylate synthases^59^ that inhibit replication of RNA viruses and retrotransposons (e.g. OAS2, > 50-fold), and prominent members of the APOBEC family of cytidine deaminases^60^ that are potent inhibitors of retroviral replication (e.g. APOBEC3G > 300-fold). When we compared the expression of these interferon-induced genes across cell types (Extended data Fig. 7c), we noted that these factors were induced in *LBR* KO, *LBR/TMPO* DKO and *LBR/TMPO/LMNA* TKO cells. Interestingly, LAP2 mRNA expression was downregulated about threefold in the *LBR* KO cells, suggesting that LBR deletion may influence the levels of both tethers.

Induction of innate immunity is not only linked to viral infection but can be caused by diverse intracellular problems, including the dysregulation of epigenetic gene silencing that is associated with the expression of retrotransposons such as endogenous retroviruses (ERVs) and inverted Alu repeats^61–63^, defects in pre-mRNA splicing ^64^, or loss of R-loop resolution that leads to the accumulation of R-loop-derived cytoplasmic RNA–DNA hybrids^65^. To deduce whether transposable elements (TEs) were deregulated upon loss of NE-heterochromatin tethers, we analysed bulk RNA-seq data for deregulation of TEs using TETranscripts^66^. This revealed a deregulation of TEs upon perturbation of heterochromatin tethering in both the *LBR/TMPO* DKO and the *LBR/TMPO/LMNA* TKO cell lines, especially of LTR retrotransposons (Fig. 5c). Notably, we also observed a dysregulation of splicing, including changes in intron retention (Extended data Fig. 7d,e).

### Loss of LBR and LAP2 affects the differentiation of mouse embryonic stem cells

LADs are known to be established shortly after fertilization^67^ and Lamin B1-DamID experiments have defined the genomic regions that are part of LADs in mESCs. The patterning of LADs is largely robust during differentiation, with cumulative changes accompanying lineage commitment and cell fate determination^57,68–71^. Interestingly, the loss of all nuclear lamins (i.e. *Lmnb1*, *Lmnb2*, *Lmna* TKO) is known to affect the global organization of chromatin in mESCs ^72^, pointing at an important function of the nuclear periphery in genome organization also in pluripotent cells.

To investigate the impact of the identified NE–heterochromatin tethers in mESCs and on differentiation, we generated *Lbr/Lmna* DKO, *Lbr/Tmpo* DKO, and *Lbr/Tmpo/Lmna* TKO cells (Extended Data Fig. 8) in the background of the E14TG2a cells, a blastocyst-derived mESC line. We maintained these cells in a naïve pluripotent ground state in presence of LIF/2i^73^. Immunofluorescence analysis of H3K9me3, H4K20me3 and H3K27me3 revealed that both *Lbr/Tmpo* DKO and *Lbr/Tmpo/Lmna* TKO cells lost the enrichment of constitutive heterochromatin at the nuclear periphery and showed an accumulation of chromocenter-like heterochromatin clusters marked with H3K9me3 and H4K20me3 in the nuclear interior (Fig. 6a-c). In contrast, *Lbr/Lmna* DKO cells maintained a large fraction of heterochromatin at the nuclear rim, while showing larger intranuclear clusters of collapsed chromatin.

**Figure 6.**
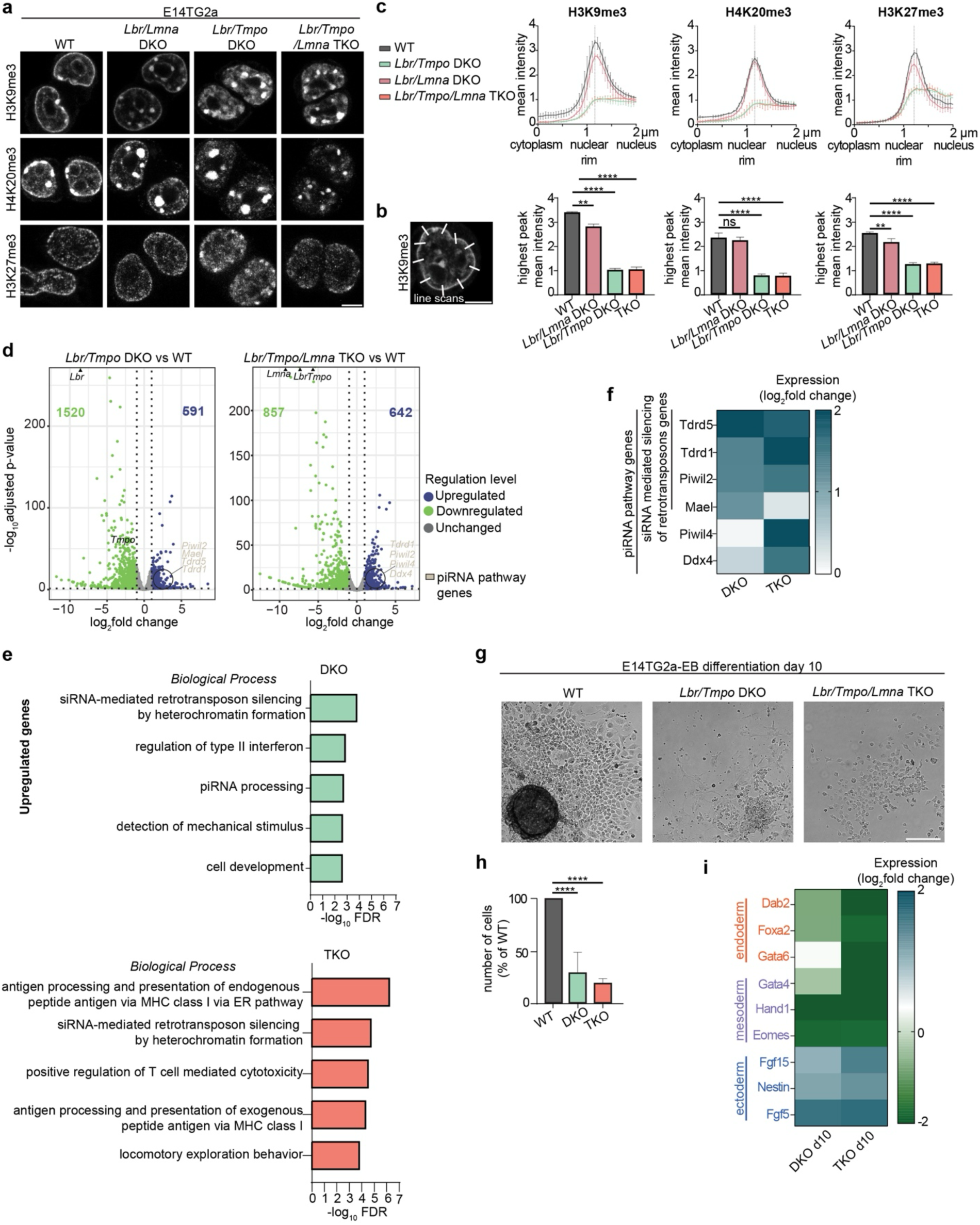
Deletion of LBR and LAP2 in mouse embryonic stem cells causes heterochromatin positioning defects, deregulation of gene expression, and aberrant differentiation. **a**, Representative confocal images of WT, *Lbr/Lmna* DKO, Lbr*/Tmpo* DKO, and *Lbr/Tmpo/Lmna* TKO E14TG2a mESCs, fixed and subjected to immunofluorescence using anti-H3K9me3, anti-H4K20me3 and anti-H3K27me3 antibodies. Note that mouse cells possess chromocenters, which are clusters of centromeric heterochromatin typically found attached to the NE in WT cells. Scale bar, 5 μm. **b**, Schematic depiction of quantification method. The fraction of antibody-stained heterochromatin at the nuclear rim was quantified using 10 line scans across the NE per analysed cell nucleus, with each line reaching 1 μm into the nucleus and the cytoplasm. **c**, Quantification of fluorescence signal at the nuclear rim in (A), plotting the normalized fluorescence intensity at each point of the line scan and the highest peak mean intensity (bar plots). *Lbr/Lmna* DKO, *Lbr/Tmpo* DKO, and *Lbr/Tmpo/Lmna* TKO mESCs (n = 12, *p ≤ 0.05, **p ≤ 0.01, ***p ≤ 0.001, ****p≤ 0.0001, Welch’s t-test, N = 3). Error bars, SD. **d**, Volcano plots of differential gene expression identified by RNA-seq (E14TG2a *Lbr/Tmpo* DKO and *Lbr/Tmpo/Lmna* TKO vs. WT cells, N = 3). Genes with an adjusted p-value ≤ 0.05 are marked as up- or downregulated if log_2_-fold change ≥ l1l. **e**, Bar graph of GO term analysis (‘Biological process’ category) of significantly upregulated genes with a log_2_-fold change ≥ 1 in *Lbr/Tmpo* DKO and *Lbr/Tmpo/Lmna* TKO cells for the category ‘Biological process’. Fischer’s exact test was used to determine overrepresented terms. **f**, Heatmap of piRNA pathway genes upregulated in *Lbr/Tmpo* DKO and *Lbr/Tmpo/Lmna* TKO cells. **g**, Representative phase contrast images of E14TG2a WT, *Lbr/Tmpo* DKO, *Lbr/Lmna* DKO, and *Lbr/Tmpo/Lmna* TKO cells after 10 days of embryoid body differentiation. Scale bar, 200 μm. **h**, Quantification of cell numbers after 10 days of embryoid body differentiation. **i**, Heatmap displaying the expression of marker genes for the three germ layers in *Lbr/Tmpo* DKO and *Lbr/Tmpo/Lmna* TKO cells as identified by RNA-seq after 10 days of embryoid body differentiation.

To examine the effects of loss of heterochromatin tethering to the NE on genome organization and gene expression, we performed ATAC-seq and RNA-seq, giving focus on *Lbr/Tmpo* DKO and *Lbr/Tmpo/Lmna* TKO cells. Compared to what we had observed in HCT116 cells (Fig. 4), ATAC-seq showed fewer changes in chromatin accessibility in mESCs (Extended Data Fig. 9a,b; Extended Data Table 1), with the larger fraction of genomic regions becoming less accessible, likely because chromatin in stem cells is generally more accessible at the outset. Still, RNA-seq identified prominent changes in gene expression (Fig. 6d), with larger fractions of genes being downregulated in the DKO and TKO cells (Extended Data Fig. 9c), consistent with the trend in the ATAC-seq data. Genes originally positioned in LADs and inter-LADs^74^ were similarly affected (Extended Data Fig. 9c).

Interestingly, GO term analysis highlighted the upregulation of groups of genes related to retrotransposon silencing, interferon response, piRNA processing, and for the TKO, adaptive immunity (Fig. 6e). The cluster related to retrotransposon silencing and piRNA processing comprised factors involved in piRNA biogenesis and function, including the RNA helicase Ddx4/Vasa, the PIWI proteins Piwil2 and Piwil4, Maelstrom (Mael) and the Tudor domain-containing proteins Tdrd1 and Tdrd5^75^ (Fig. 6f). Notably, we did not observe a significant emergence of the expression of TEs in the RNA-seq data of the mESCs (Extended Data Fig. 9d). Since both the DKO and TKO cell lines maintain expression of pluripotency markers (Pou5f1 (Oct4), Nanog) and do not show signs of precocious differentiation when maintained in LIF/2i (Extended Data Fig. 9e), it will be exciting to decipher whether the upregulation of piRNA biogenesis genes reflects an induced innate immune response involving piRNAs to the activation of retrotransposons^76^ or the emergence of a fraction of cells that have activated the piRNA pathway because they have adopted a different cell state^77^.

Finally, we used embryoid body differentiation (EB) to examine the effect of loss of peripheral heterochromatin tethering on differentiation. By withdrawal of LIF/2i, cells were induced to differentiate into the three different germ layers endoderm, mesoderm and ectoderm, and, by growth in suspension for 6 days, cells allowed to cluster forming EBs. After EB reattachment to gelatin-coated plates, we observed marked defects in both the apparent size of EBs and the total number of cells recovered after 10 days of differentiation, both for the DKO and TKO cells. These data suggest that a large proportion of cells is unable to successfully transition from pluripotency to a differentiated state (Fig. 6g,h). For the fraction of cells that successfully developed EBs, RNA-seq revealed changes in the relative abundance of markers for the three germ layers (Fig. 6i, Extended Data Fig. 9e,f; Extended Data Table 1). The expression of endoderm (Gata6, Foxa2, Dab2) and mesoderm (Gata4, Hand1, Eomes) markers was reduced in the KO cells, whereas expression of ectoderm markers (Fgf15, Nestin and Fgf5) was significantly increased, indicating that the induced changes in chromatin organization impair cell fate determination. Moreover, we noticed a strong upregulation of genes expressed in neuroectoderm, in particular of factors involved in GABAergic synaptic transmission, indicating that cells might have precociously further differentiated (Extended Data Fig. 9f). In conclusion, mESCs that have lost NE-heterochromatin tethers change their gene expression profile, have severe defects in forming EBs, and are compromised in the fidelity of cell fate determination.

## Discussion

In this study, we employed a reverse genetics approach to identify factors involved in the attachment of constitutive heterochromatin to the NE of mammalian cells. We demonstrated that the collective downregulation of a set of abundant and ubiquitously expressed NE proteins leads to the detachment of heterochromatin from the nuclear periphery. Among these factors, we identified LBR and the membrane-bound isoforms of LAP2 as the major tethers for bulk H3K9me3-marked heterochromatin including centromeres in all analysed cell types. Loss of these tethers causes changes in gene expression, chromatin organization, histone H3 methylation as well as an activation of innate immunity pathways. mESCs lacking these tethers have defects in the transition from pluripotency to differentiation and cell fate determination (Extended Data Figure 10).

### Tethering of heterochromatin to the NE determines the ‘conventional’ nuclear architecture

The current model of nuclear organization posits that the conventional radial positioning of chromatin, as existent in most differentiated mammalian cells, is driven by the active tethering of heterochromatin to the nuclear lamina^78^. This view is based on groundbreaking studies of the inverted chromatin architecture in rod cell nuclei of nocturnal animals^8^ and further supported by modelling approaches exploiting polymer simulations^9^. In agreement with this model, we show that the collective downregulation of abundant components of the nuclear lamina or the combined abrogation of the identified tethers LBR and LAP2 led to a detachment of bulk heterochromatin from the NE, providing direct experimental evidence for the importance of active tethering for peripheral heterochromatin positioning.

Upon loss of NE-heterochromatin tethering, the detached heterochromatin accumulated prominently around nucleoli. The observed relocalisation of heterochromatin to the nucleolar surface suggests that the affected genomic regions are converted from LADs into NADs^79,80^, thereby rendering nucleoli-associated heterochromatin more prominent. The observed repositioning of heterochromatin into the vicinity of nucleoli is in line with the observation that even in unperturbed, dividing cells, many genes can dynamically reshuffle between LADs and NADs^20,21^, the composition of which overlaps substantially^5,21^. Since both LADs and NADs provide a mostly repressive environment for gene expression, the observed heterochromatin relocalisation may not cause major effects on gene expression and genome homeostasis of affected loci.

In addition to the accumulation of heterochromatin around nucleoli, some detached heterochromatin formed DNA-dense foci, reminiscent of chromocenters observed in cells of rodents, plants, and insects. Although it remains to be determined whether the heterochromatin foci that we observe are *bona fide* chromocenters, their appearance upon loss of tethering lends support to a model suggesting that whether or not heterochromatin foci are formed is governed by a tug-of-war between adhesive forces executed by NE-heterochromatin tethers and cohesive forces generated by heterochromatin bundling factors^81^. Clearly, differences must exist between human and mouse cells regarding the respective prevalence of adhesion and cohesion. Since the identified NE tethers are ubiquitously expressed in both mouse and human cells, the variations in chromocenter formation between species may either arise from differences in the presence of factors involved in centromere clustering^82^ or from variances in the relative expression levels of NE tethers.

### The INM proteins LBR and LAP2 anchor H3K9me3-positive heterochromatin to the NE

Our systematic deconvolution experiments revealed a major role of the INM proteins LBR and LAP2 in the peripheral positioning of heterochromatin. Their combined contribution was observed in all analysed cell types, suggesting a generic role of the two proteins in anchoring heterochromatin in mammals. To attach constitutive heterochromatin to the nuclear periphery, the identified NE-associated tethers must be able to recognize specific molecular cues. These may include epigenetic histone marks, heterochromatin-resident proteins, or methylated DNA as in methylated CpG dinucleotides. Confirming its established role in heterochromatin tethering to the nuclear periphery^8,83,84^, loss of LBR alone already significantly affected intranuclear heterochromatin distribution. The short nucleoplasmic domain of LBR binds HP1 that in turn binds H3K9me3^85,86^, the methyl CpG binding protein MeCP2^87^, and H4K20me2^88^. Furthermore, multimerization of LBR may contribute to chromatin compaction^88^. Thus, LBR may not only serve as a heterochromatin tether, but also directly influence chromatin organization.

We identified the membrane-bound isoforms of the INM protein LAP2 as a second group of factors that promotes the attachment of constitutive heterochromatin to the NE. Currently it is unclear whether all membrane-bound isoforms contribute according to their relative expression levels or whether this function is fulfilled by any specific isoform. All LAP2 isoforms share an N-terminal DNA binding LEM-like domain of unknown specificity, followed by a BAF-binding LEM domain^89^. Since BAF interacts with DNA non-specifically^90^, the LEM domain is unlikely to confer specificity for heterochromatin. LAP2β, the longest membrane-bound isoform of LAP2, possesses an extra domain in its nucleoplasmic region (residues 219-328) that interacts with the histone deacetylase 3 (HDAC3)^91^. Together with HDAC3, LAP2β supports the peripheral anchorage of so-called lamina-associated sequences (LASs)^43^, dispersed DNA elements within LADs characterized by extended GAGA motifs. These are recognized by the transcriptional repressor cKrox, which associates with both LAP2β and HDAC3, thereby anchoring and repressing these genes at the nuclear periphery^43^. Our data indicate that LAP2’s role in tethering heterochromatin goes beyond LASs, since its downregulation affected heterochromatin in general, including centromeres. The relationship between constitutive heterochromatin marked by H3K9me3 and LASs is not yet well defined, especially since GAGA motifs are found prominently in polycomb response elements^92^.

### The A-type tether and contribution of other INM proteins

The position of LAP2 within the confines of the binary view of A- and B-type tether is presently unclear. LAP2 has originally been identified as a lamin B1 binding partner^93^, which would place it in the B-type category, but perhaps it is too simplistic to group the identified tethers according to their binding preference for A-type and B-type lamins. Interestingly, the loss of LAP2 had a greater effect on peripheral heterochromatin in the *LMNA* KO cells, and heterochromatin positioning was more strongly affected in the TKO HCT116 cells than in the *LBR/TMPO* DKO, indicating that factors other than LBR and LAP2 might contribute. These factors may include LAP1, which interact with both A-type and B-type lamins, and for which we identified a contribution to centromere localization, but also LEM domain proteins like emerin, which is known to mislocalise to the peripheral ER upon deletion of A-type lamins^94^, or the soluble, A-type lamin associated factor PRR14^95,96^ could be involved. PRR14 interacts with HP1 and depletion of PRR14 mildly affects the peripheral positioning of H3K9me3. Since the LAP2β interactome comprises PRR14 ^97^, it is worth considering a role of PRR14 in heterochromatin tethering both in the context of A-type lamins and LAP2β.

Notably, LAP2 is still expressed in the inverted rod cell nuclei of nocturnal mammals^8,44^, where both the A- and B-type tethers are functionally absent. Considering how specialized and evolutionarily fine-tuned rod cells are, there could be other mechanisms such as posttranslational modifications at play that render LAP2 in these cells incapable of heterochromatin anchorage.

### Persistent loss of heterochromatin attachment to the nuclear periphery affects genome organization, gene expression, and differentiation

To assess the importance of heterochromatin anchorage to the NE in mammalian cells, we employed a multi-omics approach to examine changes in genome organization and function after the detachment of peripheral heterochromatin. Surprisingly, short-term depletion of a suite of NE factors involved in heterochromatin tethering caused a global release of heterochromatin from the NE but had only subtle effects on gene expression. In contrast, the *LBR/TMPO* DKO and *LBR/TMPO/LMNA* TKO HCT116 cells exhibited massive changes in the transcriptome, with thousands of genes being deregulated. Perhaps cells must undergo cycles of cell divisions and DNA replication to allow changes in in the epigenome to manifest, which are then converted into changes in gene expression.

For most of the genes deregulated in the KO cells, expression was increased, irrespective of whether genes were originally positioned in LADs and inter-LADs. The massive changes in gene expression are best explained by the observed decrease in epigenetic heterochromatin marks, especially H3K27me3 and H3K9me2. The decrease in H3K27me3 in the HCT116 DKO and TKO cells suggest that H3K27me3- marked facultative heterochromatin might be especially vulnerable to derepression. In contrast, the levels of the constitutive heterochromatin marks H3K9me3 and H4K20me3 remained by-and-large unaffected. The formation of intranuclear heterochromatin clusters or repositioning into NADs may protect constitutive heterochromatin from processes that induce histone demethylation. Facultative heterochromatin, in contrast might reshuffle from the NE into the nucleoplasm, bringing it into a different, permissive chromatin environment and making it vulnerable for activation. The restoration rate of H3K27me3 after replication is highly correlated with dense chromatin^98^. Thus, genome reorganization may perturb the nucleation and spread of polycomb domains that occur every cell cycle^99^, leading to the erasure of H3K27me-marked nucleosomes over time. Our data may indicate that the peripheral positioning of polycomb domains in close vicinity to constitutive heterochromatin may support the maintenance of their epigenetic state.

Among the upregulated factors, we identified genes related to a type I interferon response in the HCT116 KO cells. This interferon response also coincided with the deregulation of LTR retrotransposons including endogenous retroviruses, indicating that loss of heterochromatin tethering and associated alterations in the epigenome may lead to their activation. In addition, we also noted changes in splicing, which may be caused by the spatial repositioning of genomic loci relative to nuclear speckles, a proposition motivated by recent correlations of gene expression and speckle proximity^100^. Currently, we do not know what the primary cause for activation of innate immunity is.

A characteristic feature of pluripotent cells is an open state of chromatin organization, enabling plasticity for reorganisation during cell fate determination^101^. The correct establishment and maintenance of different heterochromatin types accompany cell lineage determination and differentiation^57,102^. Consistent with this notion, we observed defects in cell fate determination and differentiation when NE– heterochromatin tethers were ablated in mESCs. Notably, the KO cell lines already displayed changes in gene expression in the pluripotent state. Downregulated genes prevailed in the KO mESCs, in agreement with changes observed in promoter accessibility. Among the upregulated genes, factors involved in retrotransposon silencing were prominent in GO term analysis, likely efficiently counteracting the activation of TEs. While the level of derepression of TE may differ between HCT116 and mESCs, both cell types show gene expression signatures that reflect a cellular response to TE derepression. Upon induction of differentiation, the number of mESCs forming EBs was strikingly reduced in the KO lines, indicating that cells were unable to successfully take the transition out of pluripotency and suggesting an important contribution of peripheral heterochromatin localization during a developmental period that involves significant alterations in genome organization.

We suspect that the peripheral attachment of heterochromatin will also affect other cellular features, whether it be fundamental processes such as transcription and replication or the mechanical rigidity of cell nuclei, especially in cell types that experience deformation or mechanical strain. The conventional nuclear architecture with a sheet of heterochromatin spread out underneath the NE seems to have evolved to prevent the undesired collapse and clustering of heterochromatin which may impair cellular plasticity, likely with fatal consequences for development and differentiation.

## Methods

### Antibodies

The following primary antibodies were used in this study: rabbit anti-H3K9me3 (Abcam 8898), mouse anti-H3K9me2 (Abcam 1220), rabbit anti-H4K20me3 (Abcam 9053), rabbit anti-H3K27me3 (Millipore 07-449 and Active Motif AB_2561020), rabbit anti-H3 (Abcam 1791), rabbit anti-LBR(hu) (Abcam 32535), rabbit anti-Lamin A/C (Proteintech, 10298-1-AP), mouse anti-Lamin A/C (Kerafast, LASS2D9), rabbit anti-Lamin B1 (Abcam 16048), rabbit anti Lamin B2 (Abcam 155319), rabbit anti-LAP1 (Abcam 86307), mouse anti-LAP2 (BD Transduction Laboratories 611000), rabbit anti-Lap2 (Proteintech 1465-1-AP), rabbit anti-Emerin (Abcam 40688), rabbit anti-LEMD2 (Sigma HPA017340), rabbit anti-SUN1^103^, rabbit anti-SUN2 (Sigma HPA001209), mouse anti-beta-actin (SantaCruz sc-47778), mouse anti-CLIMP-63 (Enzo life sciences ALX804-604-C100), human anti-CREST (ImmunoVision HCT0100), rabbit anti-Lbr(mouse) (Abnova PAB15583), and rabbit anti-Emerin(mouse) (Novus Biologicals NBP1-87692). The specificity of the H3K9me2 and H3K9me3 antibodies used in this study were previously validated^104^ and confirmed using siPOOLs targeting their cognate histone methyltransferases.

Secondary antibodies included: mouse Alexa Fluor 488 (ThermoFisher Scientific, A-11001), mouse Alexa Fluor 594 (ThermoFisher Scientific, A-11005), rabbit Alexa Fluor 488 (ThermoFisher Scientific, A-11034), rabbit Alexa Fluor 594 (ThermoFisher Scientific, A-11012), human Alexa Fluor 647 (ThermoFisher Scientific, A-21445), mouse Alexa Fluor Plus 680 (ThermoFisher Scientific, A32729), and rabbit Alexa Fluor Plus 800 (ThermoFisher Scientific, A32735).

### Cell lines and cell culture

HCT116 (CCL-247; ATCC), hTERT-RPE-1 (kind gift from J. Mansfeld, Institute for Cancer Research, London), and NIH/3T3 cells (kind gift from V. Vogel, ETHZ) were grown in Dulbecco’s Modified Eagle’s medium (DMEM; Gibco), supplemented with 10% fetal calf serum (FCS; Eurobio Scientific) and 100 µg/mL penicillin/streptomycin (Corning) at 37°C, 5% CO_2_. ES-E14TG2a mESCs (CRL-1821; ATCC) were grown in Dulbecco’s Modified Eagle’s medium (DMEM; Sigma) supplemented with 13% fetal calf serum (FCS; Sigma), 0.32% 2-mercaptoethanol (Gibco), and 100 µg/mL penicillin/streptomycin (Corning). Media was freshly supplemented with 1000 units/mL recombinant mouse LIF protein (ESGRO®), 1 µM of the MEK/ERK inhibitor PD0325901 (Sigma), and 3 µM of the GSK inhibitor CHIR99021 (Sigma) (2i). Cells were plated at a density of 2.65 x 10^5^ cells per cm^2^ on plates coated with 0.2% gelatin from porcine skin (Sigma). Cells were grown at 37°C, 8% CO_2_, and passaged every 48 hours using 0.05% Trypsin-EDTA for detachment (Gibco). Cells were frozen in fetal calf serum (FCS; Sigma) supplemented with 10% DMSO (Gibco).

### Molecular cloning

The coding region of full-length LAP2β was amplified from pEGFP-N3-LAP2β^105^ and cloned into KpnI and NotI sites of pCDNA5/FRT/TO (Invitrogen), which had been modified by inserting the EGFP coding sequence into the XhoI and ApaI restriction sites, allowing for the expression LAP2β-EGFP.

### Transient transfection of cells

Transfection of HCT116 cells with plasmid vectors was done using jetPRIME (Polypus). Per well of a 6-well plate,100 µL of jetPRIME buffer, 3 µL of jetPRIME and 1 µg of plasmid DNA were mixed and incubated for 15 min. This mixture was added to cells and the medium was exchanged the following day.

### RNA interference

RNAi-mediated depletion of target proteins was accomplished using siPOOLs (siTOOLs Biotech) at a working concentration of 2 nM to deplete the mRNAs encoding for the following proteins: human *LBR, LMNA, TOR1AIP1* (LAP1)*, EMD, TMPO* (LAP2)*, LEMD2, LEMD3* (MAN1)*, SUN1, SUN2, LMNB1, LMNB2, TMPO* isoform alpha*, TMPO* isoforms beta, delta and gamma, mouse *Lbr, Lmna, Tor1aip1, Emd, Tmpo, Lemd2, Lemd3, Sun1, Sun2, Lmnb1,* and *Lmnb2*. The siPOOL negative control (siTOOLs Biotech) was used for all cell lines except for HCT116 cells, for which the AllStars negative control (QIAGEN), a single non-targeting siRNA, was used instead (because high concentrations of the siPOOL control affected HCT116 cell nuclear morphology, whereas the other cell lines were seemingly unaffected). Per well of a 6-well plate, for every 1 nM of siPOOL, 20 µL of Opti-MEM (Gibco) and 0.1 µL of RNAiMAX (ThermoFisher Scientific) were mixed, incubated for 15 min at room temperature (RT) and added to cells. For mouse cells, a double pulse of RNAi was done, re-transfecting cells 48 h after the initial treatment. Samples were harvested after a total of 96 h.

### CRISPR-Cas9 mediated generation of knock-out (KO) cell lines

CRISPR RNA guides were designed using the webtools “E-CRISP” and “CHOPCHOP”. The guides 5’-*GTGAAGTGGTAAGAGGTCGA*-3’ and 5’-*CAATGTGACGCTGCCGGCCG*-3’ were used to target the first exons of the *LBR* and *TMPO* genes respectively. The DNA templates of the guide RNA along with their reverse complement strands were ordered as single-stranded DNA oligonucleotides (Merck) and annealed, yielding dsDNA inserts with overhangs complementary to the vector pC2P-Cas9^106^. The inserts were ligated into the linearized vector. To target the *LMNA* gene, the previously described pC2P-gLMNA/C vector was used^46^. 1 µg of the resulting plasmids were transfected into HCT116 cells using jetPrime (Polyplus). After 24 h, cells were detached and transferred to a 10 cm plate and selected with 10 µg/mL of puromycin (Sigma) for 48 h. Puromycin was washed away, and clones resistant to puromycin were picked, expanded and characterised. Clones containing frameshift mutations were identified using PCR and sanger sequencing, with indels being identified using the tide web tool^107^ and through manual inspection of the sequencing chromatograms. To screen the *LBR* KO clones via PCR, the primers 5’-*GATCACGGTAGGTGGGTAACA*-3’ and 5’-*ATCCTCTGCCTTCAAACCGA*-3’ were used. The *LBR* KO clone #2 was used for the creation of the double and triple KO cell lines and in the corresponding RNA-seq experiments To screen the KO clones for edits in the *TMPO* gene, the primers 5’-*TTTGTGTCCGCGAGTTTTTGTTCCG*-3’ and 5’-*GCCAAACGCCCGCCTTTGTAG*-3’ were utilized. Finally, edits made in the *LMNA* gene were uncovered using primers 5’-*TTTCTCCCCGACCCTTTTGC*-3’ and 5’- *GGCAAAGTTATCGGCCTCCA*-3’.

CRISPR-Cas9-mediated gene knockout of *Lbr*, *Tmpo* and *Lmna* in ES-E14TG2a cells was done as previously described^108^. In brief, paired CRISPR/Cas9 sgRNAs targeting the start and stop codons of the gene of interest were designed. The guide pair 5’-*GTTGAGGGTGAAGTGGTCAG*-3’ and 5’-*GTAGCACGGCATGATCAGAC*-3’ was used to target *Lbr*, 5’-*CGGGCACGGCACAGAACTCG*-3’ and 5’-*CCAAGGGAGCATACTTCGGGACG*-3’ to target *Tmpo*, and 5’-*GCGGCGCGCCACCCGCAGTG*-3’ and 5’-*CTCGGCCTCGTGTAGTGCCG*-3’ to target *Lmna*. The respective sgRNAs targeting the start codon were cloned into pLentiCRISPR-EGFP (Adgene plasmid ID: 75159), while those targeting the stop codons were cloned into pLentiCRISPR-mCherry (Adgene plasmid ID: 75161), whereas for *Lbr*, both guides were cloned into pX458-GFP (Adgene plasmid ID: 48138). 160’000 cells were seeded into a well of a 6-well plate. For co-transfection of both vectors the next day, 2 μg of each vector was added to 250 μL of reduced-serum medium (Opti-MEM; Gibco). In parallel, 250 μL of Opti-MEM (Gibco) was mixed with 10 μL of Lipofectamine 2000 (L2000; Invitrogen). The two solutions were combined and incubated at RT for 20 min. Then, 500 μL of the mixture was added to cells to yield a final volume of 1.5 mL of medium. Media was exchanged after 6 h. 48 h after transfection, cells were sorted (BDFACSAria, Flow Cytometry Core Facility, ETHZ) to isolate single GFP and mCherry double-positive cells, or in the case of *Lbr*, GFP-positive cells. These were plated in gelatin-coated 96-wells and grown to colonies. Clones were characterized by PCR using primer pairs (PPs) flanking the sgRNAs designed with Primer-BLAST^109^, immunoblotting and by sequencing across the gene deletion site. The following PPs flanking the sgRNAs binding sites were used: 1. *Lbr* KO: PP1-F 5’-*AGTTGTCCAGTCTGATGCCTG-*3’, PP1-R 5’-*TGGGCCTATACCTCACTCCC*-3’; and PP2-F 5’-*ACAGTGTTTTTGCTACTGTTTAGC*-3’, PP2-R 5’-*ACAAAGCTTGCCAAACATCAAAT*-3’; 2. *Tmpo* KO, PP1-F 5’-*TATCAGCGTCCGAGGGGAA*-3’, PP1-R 5’-*TGCAGGTAGAGCTGCACGTA*-3’, and PP2-F 5’-*GTGATACGTCAATCCCCTGCT*-3’, PP2-R 5’-*TGAGATCCGTGTAAGCATCCTG*-3’; 3. *Lmna* KO, PP1-F 5’-*AAGGGGTGTTCTGCAGAGTG*-3’, PP1-R 5’-*AGAATCAAGGGTCTTGCGGG*-3’, and PP2-F 5’-*CTTGCAGTCCCTCTTGCATC*-3’, PP2-R 5’-*AATCGCCGCACCTCTAGACT*-3’. The E14TG2a *Lbr* KO cell line was used to generate the *Lbr/Tmpo* and the *Lbr/Lmna* DKO cell lines. The *Lbr/Lmna* DKO was used as a background for the generation of the *Lbr/Tmpo/Lmna* TKO cell line.

### Embryoid body differentiation assay

Embryoid body (EB) differentiation assays were performed as described^110^. In brief, at day zero, 1 x 10^6^ cells were seeded in 10 cm petri dishes using media lacking LIF and 2i, and carefully spread evenly across the plate by swirling. On day three, EBs were plated onto two 10 cm dishes. At day six, EBs from each dish were transferred into gelatin-coated T75 flasks and grown until day ten. At days zero and ten of differentiation, cells were analysed by phase contrast microscopy with a Nikon Ti-E with air objective Plan Fluor 10x NA 0.30 Ph1 DL WD 15.2 mm. Images were acquired with the Nikon NIS-Elements software, and samples were taken for immunoblotting and RNA extraction.

### Immunoblot analysis

Whole cell extracts were generated by taking up cells in SDS sample buffer (75 mM Tris pH 7.8, 20% (v/v) glycerol, 4% SDS, 50 mM DTT and 0.1% bromophenol blue) and separated by SDS-PAGE. Proteins were transferred onto a nitrocellulose membrane (Amersham Protran, 10600015) by semi-dry blotting. The membranes were incubated thrice in blocking solution (5% (w/v) milk powder, 0.1% Tween in 1x PBS) for 5 min. The membrane was incubated with the primary antibody diluted in blocking solution at 4°C ON, washed with blocking solution thrice for 5 min, and then exposed to fluorescently labelled secondary antibodies diluted in blocking solution for 35 min at RT. Finally, the membrane was washed thrice with 0.1% Tween in 1x PBS for 5 min. Signals were detected using an Odyssey (LI-COR) imaging system.

### Immunofluorescence

For immunofluorescence, cells on coverslips were fixed with 2% paraformaldehyde (Sigma-Aldrich) in phosphate-buffered saline (PBS) for 10 min at RT. Cells were washed twice for 3 min using 1x PBS, permeabilised with 0.2% Triton-X100 (Sigma) or 0.2% Triton-X100 and 0.02% SDS in PBS for 12 min at RT, then washed thrice with 1x PBS for 5 min. Blocking was performed in blocking solution (2% bovine serum albumin (BSA, Pan-Biotech)) and 0.1% Tween in 1x PBS) for at least 45 min at RT. Primary antibodies were incubated O/N at 4°C in blocking solution at different concentrations (anti-H3K9me3 1:500, anti-H3K27me3 1:500, anti-H4K20me3 1:500, anti-LAP2 1:2000, anti-LBR 1:1000, anti-Lamin A/C 1:1000, anti-CREST 1:1000, anti-CLIMP63 1:200). The following day, coverslips were washed thrice for 3 min using washing buffer (0.1% Tween in 1x PBS). Secondary antibodies were diluted 1:300 in blocking solution, added to the coverslips and incubated for 35 min at RT. Then, coverslips were washed thrice using washing buffer, before a final wash with 1x PBS for 3 min. Lastly, the cells were stained with 1 µg/mL Hoechst (Sigma-Aldrich, 63493) for 10 min at RT and washed with 1x PBS. The coverslips were mounted onto glass slides using 3 µL of VECTASHIELD (Vector Laboratories) and sealed with nail polish. Samples were imaged with a Zeiss LSM 880 or a Zeiss LSM 780 Confocal Laser Scanning Microscope, using an 63x 1.4 NA oil plan-apochromat objective.

### Quantification of fluorescently labelled chromatin at the nuclear periphery

To quantify the fraction of antibody-labelled heterochromatin at the nuclear periphery in differentiated cells, cells were subjected to immunofluorescence with the indicated antibodies. Images were first split into separate channels. Segmentation of cell nuclei was performed by exploiting the Hoechst channel using the nucleAIzer model^111^. To compute the nuclear rim, mathematical morphology was employed to erode the mask image with a rectangular structuring element. The size of the structuring element was empirically set to 17, which is equivalent to 8 pixels of erosion (∼0.42 µm). The difference between the nuclear mask and its eroded version determined the pixels that belong to the nuclear rim. If a few pixels of random shift were noted in the images, this was fixed by calculating the phase cross correlation between the channels and transforming the heterochromatin channel. The rim mask was then applied to the channel used for heterochromatin labelling and the underlying pixel intensities were summed and divided by the area of the rim mask. The same was done with the entire nucleus, and the average nuclear intensity was determined. The ratio of the average rim and nuclear intensities was used for further statistics. The segmented images were individually inspected, and cells for which the segmentation was poor or faulty, for instance two cells being counted as one, were excluded. The data was further analysed using GraphPad Prism 10 (GraphPad), differences between samples tested for significance (Welch’s T-test or one-way ANOVA), and graphs plotted.

To quantify the fraction of heterochromatin at the nuclear periphery of ES-E14TG2a cells the above method described did not produce robust results due to the variable size and presence of chromocenters at the nuclear periphery. Therefore, for these cells, quantification of the signal intensity of histone marks at the nuclear periphery was performed using Fiji ImageJ^112^. Per cell nucleus, ten line scans of 2 μm distributed along the entire nuclear perimeter were taken, each extending 1 μm into the cytoplasm and 1 μm into the nuclear interior. The intensity of each point of the line scan was normalised against the total nuclear signal and plotted using GraphPad. An ordinary one-way ANOVA test was performed on normalised values corresponding to the peak signal at the NE using GraphPad Prism 10.

### Quantification of centromere positioning

To measure the fraction of centromeres at the nuclear periphery, fixed HCT116 cells were subjected to immunofluorescence using anti-CREST and anti-CLIMP63 antibodies, stained with Hoechst and images taken on a Zeiss LSM 880 Confocal Laser Scanning Microscope, using an 63x 1.4 NA oil plan-apochromat objective, taking 33 z-stacks of 0.39 µm step size. These images were processed in Fiji, and composite stack of images containing Hoechst, a nuclear envelope marker (CLIMP-63), and centromeres (CREST) was generated. Using the measure function of Fiji, the number of centromeres positioned within a 0.5 µM-wide ring at the nuclear periphery was determined. The distance of 0.5 µM was considered as within the range of a LAD based on previous work^20,71^. Each centromere focus was only counted once, in the plane where it was the brightest. For the distance measurements, the measuring pointer was placed at the centre of each focus, and the shortest distance to the nuclear periphery was determined. If centromere foci were too close to each other not allowing to resolve single centromeres, these foci were counted as one centromere. After the peripheral centromeres had been counted and measured, the number of internal centromeres was determined to calculate the fraction of peripheral centromeres over all centromeres. GraphPad Prism 10 was used to perform statistical tests and plot graphs.

### Total RNA extraction and RNA-seq

To prepare samples for RNA-seq, 3 individual batches of cells were plated on the same day and further processed in parallel. Freshly harvested cells were lysed with 500 µL of QIAzol lysis reagent (QIAGEN #79306) and mixed by gentle pipetting. Subsequently, 100 µL of chloroform (Sigma) was added, samples were vortexed for 10 s and spun at 12’000 x g for 15 s at 4°C. The upper phase was transferred into a new tube, 600 μL of isopropanol (Sigma) added and samples incubated O/N at −20°C. After centrifugation at 16’000 x g for 10 min at 4°C, the RNA pellet was washed in 75% ethanol and resuspended in RNase-free water (Invitrogen). DNase digestion of RNA was performed using RNase-Free DNase Set (QIAGEN) following the manufacturer’s instructions. RNA was then precipitated by adding sodium acetate (Sigma) to a final concentration of 0.3 M, and adding 3 volumes of 100% ethanol O/N at −20°C. RNA was recovered by centrifugation at 16’000 x g for 10 min and washed two times with 70% ethanol. RNA was dissolved in RNase-free water (Invitrogen).

RNA sequencing was performed at the Functional Genomic Center Zurich. The quality of the isolated RNA was determined using a Fragment Analyzer (Agilent, Santa Clara, California, USA). Only samples with a 260 nm/280 nm ratio between 1.8–2.1 and a 28S/18S ratio within 1.5–2 were further used for library preparation. The Illumina Ribo Zero Plus (Illumina, Inc, California, USA) was used in the succeeding steps. Briefly, total RNA samples (25-1000 ng) were depleted of ribosomal RNA and then reverse-transcribed into double-stranded cDNA. The cDNA samples were fragmented, end-repaired, and adenylated before ligation of an anchor. Fragments containing the anchor on both ends were selectively enriched with PCR at the same time adding the index with unique dual indexes (UDI). The quality and quantity of the enriched libraries were validated using the Fragment Analyzer (Agilent, Santa Clara, California, USA). The libraries were normalized to 10 nM in Tris-HCl 10 mM, pH8.5 with 0.1% Tween20.The Novaseq X (Illumina, Inc, California, USA) was used for cluster generation and sequencing according to the standard protocol. Sequencing experiments were paired-end at 2 x 150 bp.

### ATAC-seq

To prepare samples, 3 individual batches of cells were plate on the same day in 6-wells (for si-11) or 10 cm plates (for KO cells). Cell batches were then further processed in parallel, during all subsequent steps. For harvest, cells were washed twice with 1x PBS and detached using 0.5 mM EDTA in 1x PBS. Cells were counted and 80’000 cells were pelleted. Pellets were resuspended in 200 µL of freezing medium (10% DMSO in FCS) and slowly frozen at −80°C. ATAC-seq was performed at QuickBiology (Monrovia, California). Here, cells were washed and lysed to generate a crude preparation of nuclei. This crude extract was then treated with Tn5 Transposase. Tn5 transposase tagmentation simultaneously fragments the genome and tags the resulting DNA with Illumina sequencing adapters. DNA fragments were PCR-amplified and subsequently purified using the MinElute PCR Purification Kit (Qiagen, Maryland, USA). The ATAC-seq library was prepared as described protocol^113^. After quality control using an Agilent TapeStation 4200 (Agilent Technologies, Santa Clara, CA) and Life Technologies Qubit 3.0 Fluorometer (Life Technologies, Carlsbad, CA), the libraries were sequenced (150 bp paired-end reads) on an Illumina HiSeq Sequencer (Illumina Inc., San Diego, CA).

### Analysis of multi-omics data

#### Reference genomes and annotations

The reference genomes for human (GRCh38; annotation version 108) and mouse (GRCm39; annotation version 112) were obtained from ENSEMBL^114^.

#### Bulk RNA sequencing

The paired-end reads from bulk RNA-seq datasets were processed to remove the Illumina universal adapter (*AGATCGGAAGAGCACACGTCTGAACTCCAGTCA*) using Cutadapt (v4.4) with the parameters “*-q 25 -m 25*”^115^. The resulting filtered reads were aligned to their respective reference genome using STAR aligner (v2.7.10b) with default parameters^116^. Coordinate-sorted alignment files were indexed using SAMtools (v1.17)^117^. Exon mapped reads were counted using featureCounts from the subread package (v2.0.6)^118^. Subsequent data analysis was performed with custom scripts in R (R Core Team (2024). Data wrangling and visualization was performed using tidyverse package (v2.0.0)^119^. Differential expression analysis was conducted using DESeq2 (v1.44)^120^. Differentially expressed genes (DEGs) were defined as having absolute log fold change > 1 and adjusted p-value < 0.05. DEGs were further analysed for gene ontology analysis using TopGO (Bioconductor.org)^121^.

To evaluate the expression of transposable elements, the filtered reads from above were aligned to their respective reference genome using STAR aligner (v2.7.10b) with the parameters “*--winAnchorMultimapNmax 200 --outFilterMultimapNmax 100*”. The resulting alignment files were used to quantify read counts for genes and transposable elements using TETranscripts (v2.2.3). Differential expression analysis was performed using DESeq2 (v1.44) and differentially expressed TEs were defined as having absolute log fold change > 0.585 and adjusted p-value < 0.05.

#### ATAC sequencing

ATAC-seq datasets were processed to remove the Nextera adapter (*CTGTCTCTTATACACATCT*) using Cutadapt (v4.4) with the parameters “*-q 25 -m 25*”^115^. The resulting filtered reads were aligned to reference human genome using bowtie2 aligner (v2.5.1) with the parameter “--very-sensitive” (Bowtie2^122^). The alignment file was sorted by coordinate, indexed and filtered for mitochondrial reads using SAMtools (v1.17). The filtered alignment file was further processed to keep only uniquely mapped, non-duplicated reads using sambamba (v1.0.1)^123^. Reads were further processed using *alignmentSieve* from deepTools (v3.5.5) to account for transposon dimerization before insertion. Positive and negative strand alignments were shifted by +4 bp and −5 bp, respectively. Additionally, fragments of 100 bp, corresponding to nucleosome-free regions (NFRs), were retained with the parameters “-*-minMappingQuality 25 --maxFragmentLength 100 --ATACshift --BED*”^124^. Peaks were called on the resulting BEDPE files using MACS3 (v3.0.1) MACS3^125^ with the parameters “*--call-summits --bdg --nomodel --nolambda --gsize hs --keep-dup all --format BEDPE --qvalue 0.05*”. The predicted peaks were filtered for ENCODE project blacklisted regions using BEDTools (v2.31.0) intersect (ENCODE blacklist^126^; BEDTools^127^).

Peaks from the three replicates of a condition were intersected using BEDTools (v2.31.0) with the parameter “*-f 0.50 -r*”^127^. Peaks detected in at least 2 out of 3 replicates of a condition were considered consensus regions corresponding to open chromatin. Differentially accessible regions (DARs) were identified using DESeq2 (v1.44). DARs were annotated using Chipseeker package (v1.40)^128^.

## Acknowledgements

We thank I. Zemp, M. Jagannathan, T. Kleele, H. Baird, and S. Pawar for critical comments on the manuscript, members of the Kutay lab for discussions, and M. Jagannathan, R. Foisner, A. Wutz, T. Aktas, Z. Smith and their teams for helpful suggestions. Microscopy was performed on instruments of the Microscopy Center Scope M of ETHZ, RNA-seq was supported by C. Aquino and D. Rodrigues at the Functional Genomics Center Zurich, and the processing of genomics data was performed on the Euler high-performance computing cluster at ETH Zürich.

This work was supported by the Swiss National Science Foundation (SNSF) [310030_219203 and TMAG-3_209245 (U.K.)]. We thank the Swiss NCCR ‘RNA and disease’ for networking opportunities. PH and KK acknowledge support from the Lendület BIOMAG grant [2018–342], TKP2021-EGA09, Horizon-BIALYMPH, Horizon-SYMMETRY, Horizon-SWEEPICS, H2020-Fair-CHARM, CZI Deep Visual Proteomics, HAS-NAP3, the ELKH-Excellence grant from OTKA-SNN [139455/ARRS] and Finnish Cancer Society.

## Data and code availability

High-throughput sequencing data generated in this study are deposited at NCBI gene expression omnibus (GEO) database under the accession number GSEXXXXX, and are publicly available as of the publication date.

All relevant code used for the analysis of high-throughput sequencing data is available at GitHub (XYZ).

