## Supplemental Material for "LBR and LAP2 mediate heterochromatin tethering to the nuclear periphery to preserve genome homeostasis"

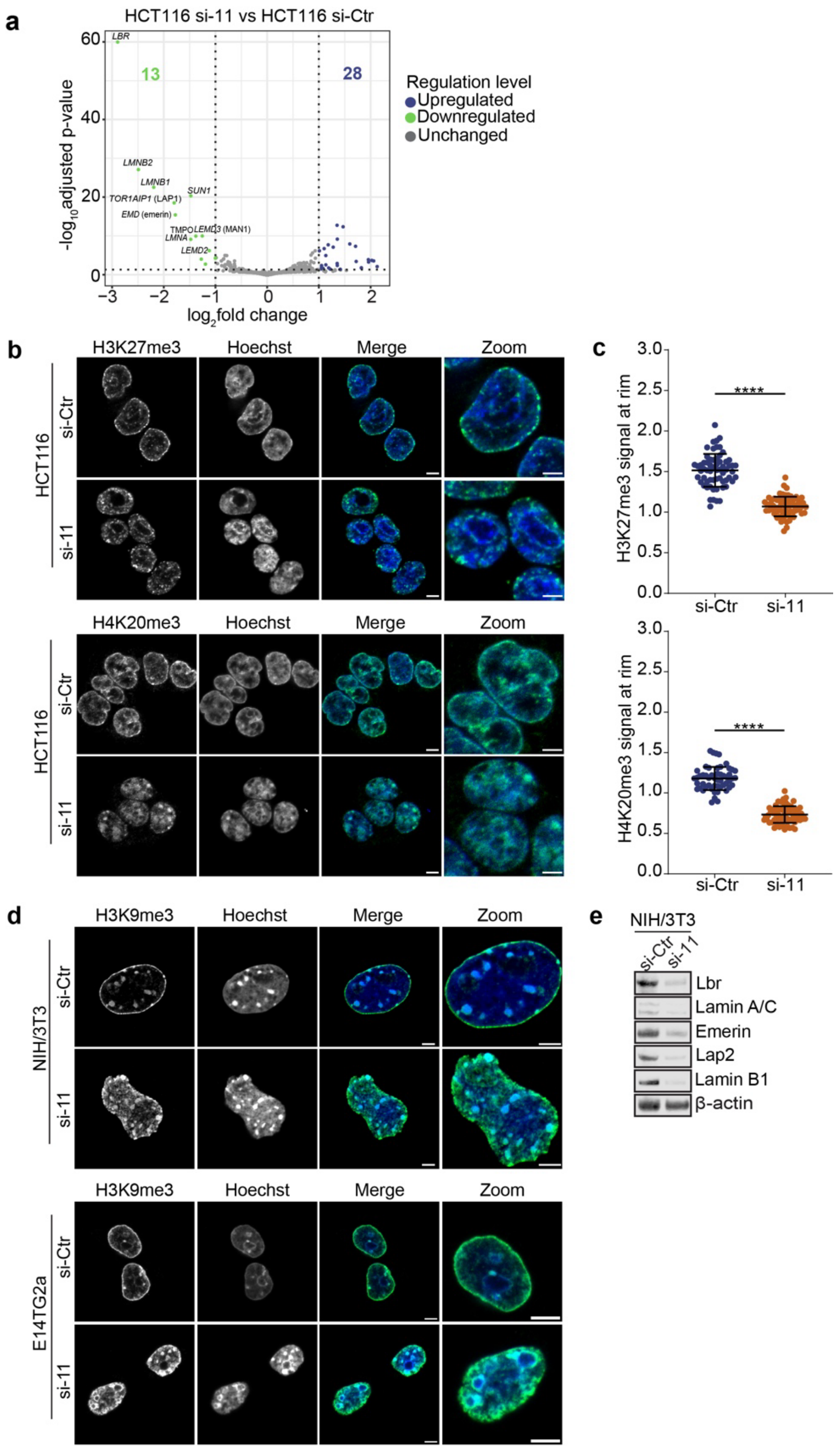


**Figure S1. H3K9me3-stained heterochromatin is detached from the nuclear periphery of mouse differentiated and pluripotent cells upon depletion of 8 INM proteins and the nuclear lamins.**

**a**, Volcano plot of si-11-treated vs. si-Ctr treated HCT116 cells (N = 3). Genes with a $\log_2$-fold change ≥ I1I and an adjusted p-value ≤ 0.05 were considered significant.

**b**, Representative confocal images of HCT116 cells that were treated either with si-Ctr or si-11 for 72 h, fixed and subjected to immunofluorescence using an anti-H3K27me3 and anti-H4K20me3 antibody. DNA was stained using Hoechst. Scale bars, 5 μm.

**c**, Quantification of H3K27me3 and H4K20me3 at the nuclear rim in HCT116 cells that were either treated with si-Ctr or si-11 (n ≥ 52, *p ≤ 0.05, **p ≤ 0.01, ***p ≤ 0.001, ****p≤ 0.0001, Welch's t-test, N= 3). Error bars, SD.

**d**, Representative confocal images of NIH/3T3 cells and mESCs (E14TG2a) treated either with control siRNAs (si-Ctr) or a combination of 11 siPOOLs (si-11) for 72 h, fixed and subjected to immunofluorescence using an anti-H3K9me3 antibody. DNA was stained using Hoechst. Scale bars, 5 μm.

**e**, Immunoblots confirming the downregulation of select mouse NE proteins.

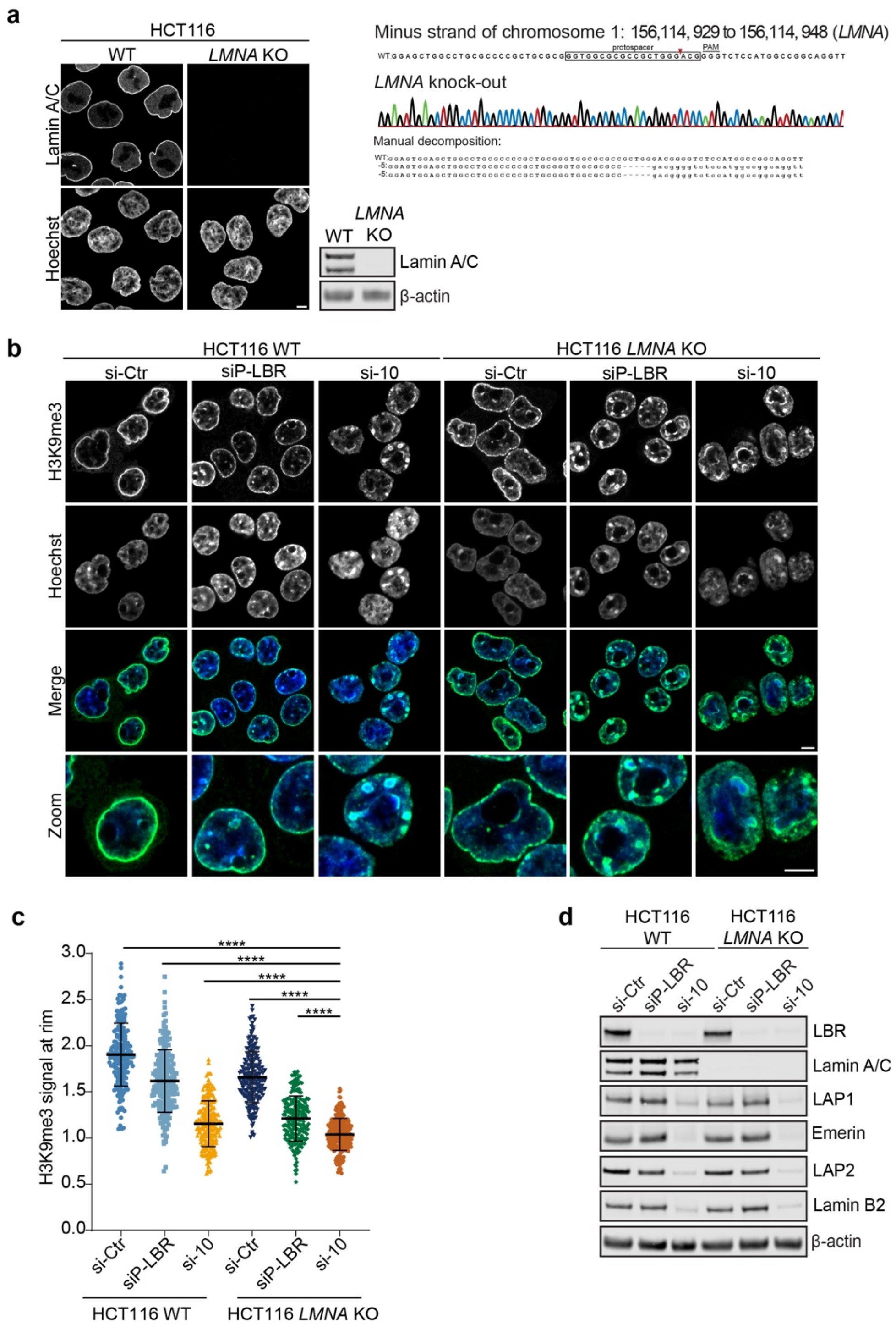


**Figure S2. H3K9me3-stained heterochromatin is detached from the nuclear periphery in HCT116 *LMNA* knockout cells upon the collective depletion of 8 INM proteins and the B-type lamins.**

**a**, Characterisation of the HCT116 *LMNA* KO clone. Immunofluorescence and immunoblotting analyses of lamin A/C in parental HCT116 WT and HCT116 *LMNA* KO using anti-lamin A/C antibodies. Manual decomposition of the sequence surrounding the PAM in the *LMNA* KO clone confirms frameshift mutations in both alleles. Genomic coordinates correspond to the location where the guide RNA binds. Scale bar, 5 µm.

**b**, Representative confocal images of HCT116 WT and HCT116 *LMNA* KO cells that were either treated with si-Ctr, an siPOOL against LBR (siP-LBR) or a combination of 10 siPOOLs (si-10), directed against lamins B1 and B2 and the INM proteins LBR, LAP1, Emerin, LAP2, LEMD2, MAN1, SUN1, and SUN2) for 72 h, fixed and subjected to immunofluorescence using an anti-H3K9me3 antibody. Scale bars, 5 µm.

**c**, Quantification of the H3K9me3 signal at the nuclear rim (n ≥ 199, *p ≤ 0.05, **p ≤ 0.01, ***p ≤ 0.001, ****p≤ 0.0001, one-way ANOVA, N = 3). Error bars, SD.

**d**, Immunoblotting of lamin A/C, LBR and select other NE proteins to determine the efficiency of siRNA-induced protein depletion.

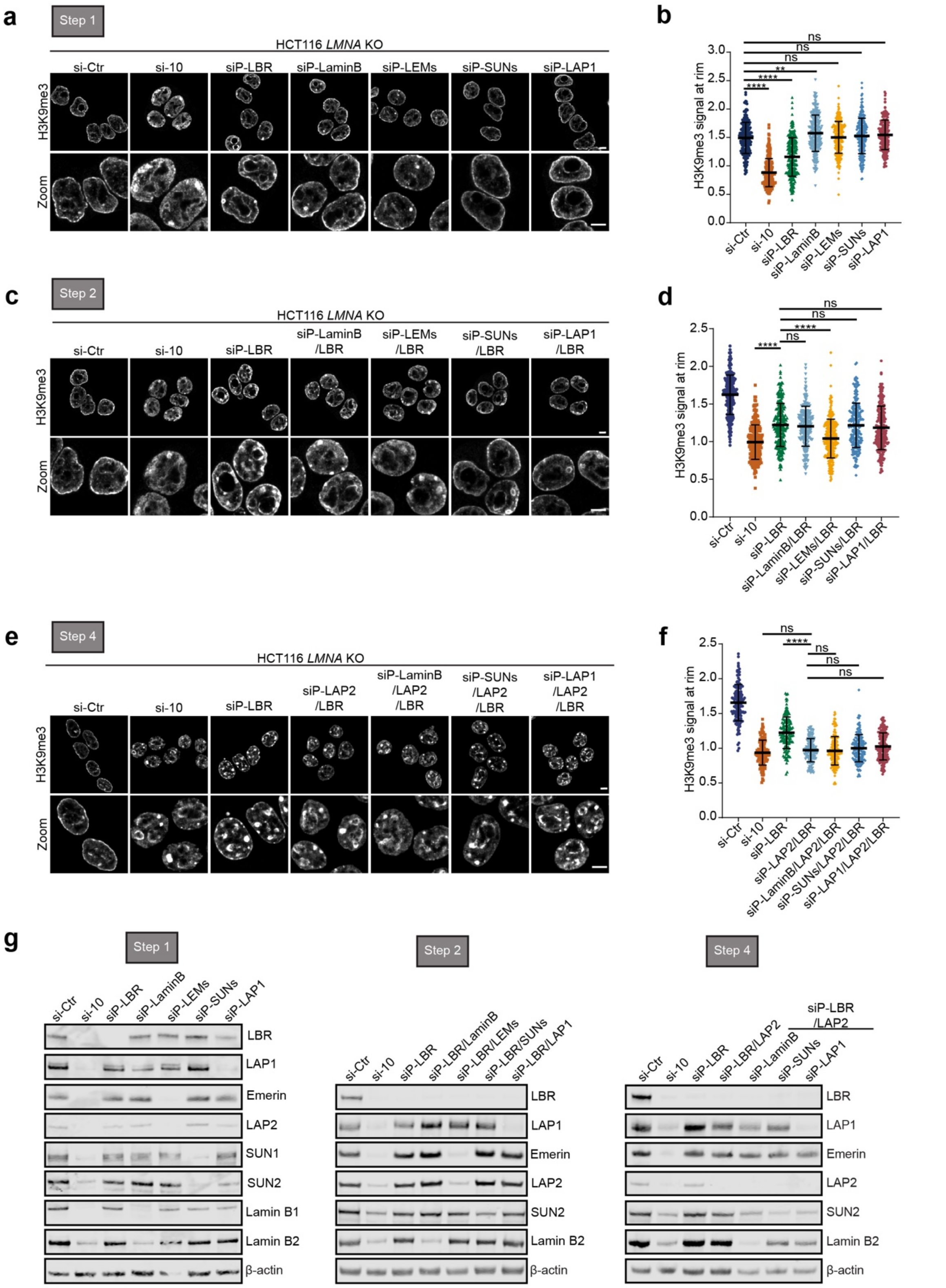


**Figure S3. The INM proteins LBR and LAP2 promote the peripheral positioning of H3K9me3-positive heterochromatin.**

**a**, Representative confocal images of H3K9me3-positive heterochromatin in HCT116 *LMNA* KO cells treated with si-Ctr or the indicated combinations of siPOOLs for 72 h (Step 1 of deconvolution), identifying LBR. Scale bars, 5 μm.

**b**, Quantification of the H3K9me3 signal at the nuclear rim ($n \geq 234$, *$p \leq 0.05$, **$p \leq 0.01$, ***$p \leq 0.001$, ****$p \leq 0.0001$, one-way ANOVA, $N = 4$). Error bars, SD.

**c**, Representative confocal images of H3K9me3-positive heterochromatin in HCT116 *LMNA* KO cells treated with si-Ctr or the indicated combination of siPOOLs for 72 h (Step 2 of deconvolution), identifying LEM-domain family of proteins. Scale bars, 5 µm.
**d**, Quantification of the H3K9me3 signal at the nuclear rim ($n \geq 215$, *$p \leq 0.05$, **$p \leq 0.01$, ***$p \leq 0.001$, ****$p \leq 0.0001$, one-way ANOVA, N = 4). Error bars, SD.
**e**, Representative confocal images of H3K9me3-positive heterochromatin in HCT116 *LMNA* KO cells treated with si-Ctr or the indicated combination of siPOOLs for 72 h (Step 4 of deconvolution). Scale bars, 5 µm.
**f**, Quantification of the H3K9me3 signal at the nuclear rim ($n \geq 142$, *$p \leq 0.05$, **$p \leq 0.01$, ***$p \leq 0.001$, ****$p \leq 0.0001$, one-way ANOVA, N = 3). Error bars, SD.
**g**, Immunoblotting of LBR and select other NE proteins to determine the efficiency of siRNA-induced protein depletion.

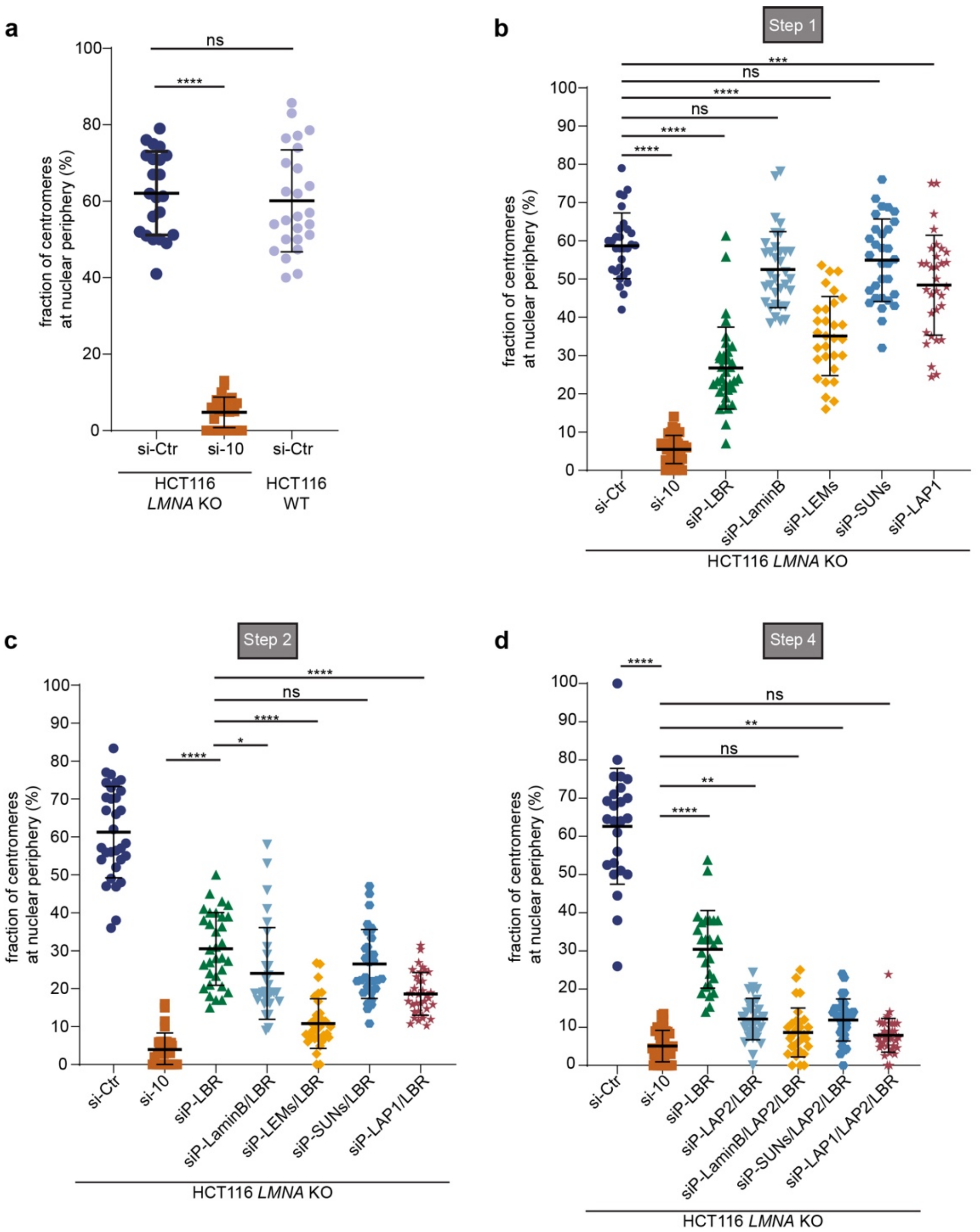


**Figure S4. The INM proteins LBR, LAP2 and LAP1 promote the peripheral positioning of centromeres.**

**a**, Quantification of a proof-of-concept experiment to measure the proportion of centromeres within in 0.5 µm-wide ring at the nuclear periphery in si-Ctr and si-10 treated HCT116 *LMNA* KO cells as well as in si-Ctr treated HCT116 WT cells (n ≥ 22, *p ≤ 0.05, **p ≤ 0.01, ***p ≤ 0.001, ****p≤ 0.0001, one-way ANOVA, N = 2). Error bars, SD.

**b**, Quantification of the proportion of centromeres at the nuclear periphery in HCT116 *LMNA* KO cells treated with si-Ctr or the indicated combination of siPOOLs for 72 h (Step 1 of deconvolution), identifying LBR (n ≥ 29, *p ≤ 0.05, **p ≤ 0.01, ***p ≤ 0.001, ****p≤ 0.0001, one-way ANOVA, N = 3). Error bars, SD.

**c**, Quantification of the proportion of centromeres at the nuclear periphery in HCT116 *LMNA* KO cells treated with si-Ctr or the indicated combination of siPOOLs for 72 h (Step 2 of deconvolution), identifying the LEM-domain family of proteins (n ≥ 30, *p ≤ 0.05, **p ≤ 0.01, ***p ≤ 0.001, ****p≤ 0.0001, one-way ANOVA, N = 3). Error bars, SD.

**d**, Quantification of the proportion of centromeres ≤ 0.5 µm at the nuclear periphery in HCT116 *LMNA* KO cells treated with si-Ctr or the indicated combination of siPOOLs for 72 h (Step 4 of deconvolution), identifying the INM protein LAP1 (n ≥ 25, *p ≤ 0.05, **p ≤ 0.01, ***p ≤ 0.001, ****p≤ 0.0001, one-way ANOVA, N = 3). Error bars, SD.

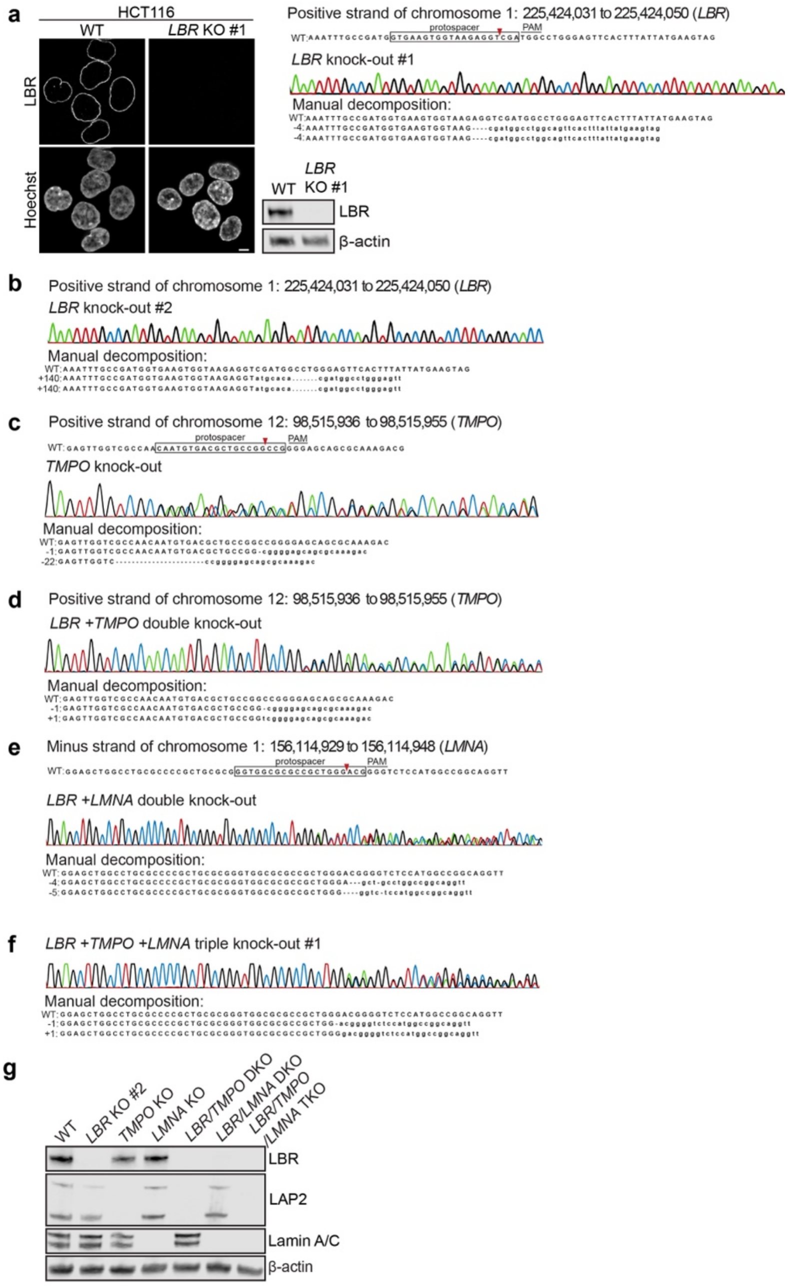


**Figure S5. Characterization of HCT116 knock-out cell lines.**

**a**, Characterization of the HCT116 *LBR* KO clone #1. Immunofluorescence and immunoblotting analyses of LBR in parental HCT116 WT and HCT116 *LBR* KO using anti-LBR antibodies. Manual decomposition of the sequence surrounding the PAM in the *LBR* KO clone #1 confirms frameshift mutations in both alleles. Genomic coordinates correspond to the location where the guide RNA binds. Used in Fig. 3, Fig. S6g and Fig. S7c.
**b**, Characterization of the HCT116 *LBR* KO clone #2. Manual decomposition of the sequence surrounding the PAM in the *LBR* KO clone #2 confirms frameshift mutations in both alleles. Used in Fig. 4, 5, Fig. S6e and Fig.S7e.
**c**, Characterization of the HCT116 *TMPO* KO clone. Manual decomposition of the sequence surrounding the PAM in the *TMPO* KO clone confirms frameshift mutations in both alleles.
**d**, Characterization of the HCT116 *LBR +TMPO* DKO clone. Manual decomposition of the sequence surrounding the PAM in the *LBR +TMPO* DKO clone confirms frameshift mutations in both alleles. Derived from LBR KO clone #2.
**e**, Characterization of the HCT116 *LBR +LMNA* DKO clone. Manual decomposition of the sequence surrounding the PAM in the *LBR +LMNA* DKO clone confirms frameshift mutations in both alleles. Derived from LBR KO clone #2.
**f**, Characterization of the HCT116 *LBR +TMPO +LMNA* TKO clone #1. Manual decomposition of the sequence surrounding the PAM in the *LBR +TMPO +LMNA* TKO clone #1 confirms frameshift mutations in both alleles. Derived from LBR KO clone #2.
**g**, Immunoblot of LBR, LAP2 and lamin A/C in various KO backgrounds to validate the loss of these proteins.

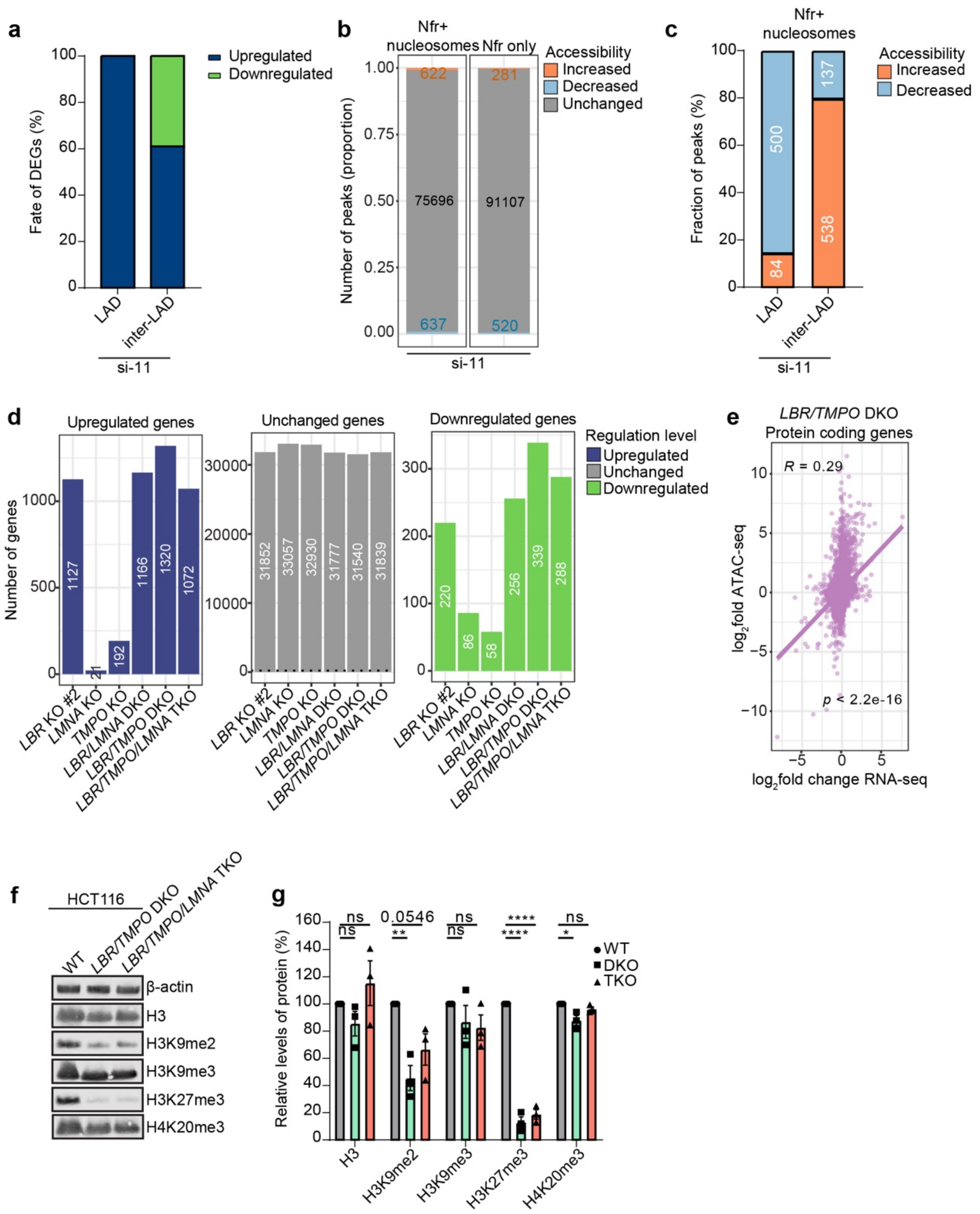


**Figure S6. Changes in the transcriptome and chromatin organization upon the loss of heterochromatin tethers.**

**a**, Bar graph showing the fate of deregulated genes previously positioned in LADs and inter-LADs in si-11 treated cells.

**b**, Proportion of Nfr nucleosomes and Nfr only ATAC-seq peaks with increased or decreased accessibility in si-11 treated cells (N = 3). All loci with a $\log_2$-fold change ≥ I1I and an adjusted p-value ≤ 0.05 were considered significant.

**c**, Proportion of ATAC-seq peaks in both Nfr plus nucleosome-containing regions with increased or decreased accessibility in regions designated as LADs or inter-LADs in si-11 treated cells.

**d**, Bar graph of differentially expressed genes identified by RNA-seq in all genotypes (N = 3). Genes with an adjusted p-value ≤ 0.05 are marked as up- or downregulated if $\log_2$-fold change ≥ I2I.

**e**, Correlation analysis between changes in RNA-seq and ATAC-seq.

**f**, Representative immunoblot of H3, H3K9me2, H3K9me3, H3K27me3 and H4K20me3 relative to β-actin in *LBR/TMPO* DKO and *LBR/TMPO/LMNA* TKO cells. N = 3.

**g**, Quantification of protein levels (normalized to β-actin) for histone modifications in *LBR/TMPO* DKO and *LBR/TMPO/LMNA* TKO cells (*p ≤ 0.05, **p ≤ 0.01, ***p ≤ 0.001, ****p≤ 0.0001, one-way ANOVA, N = 3). Error bars, SEM.

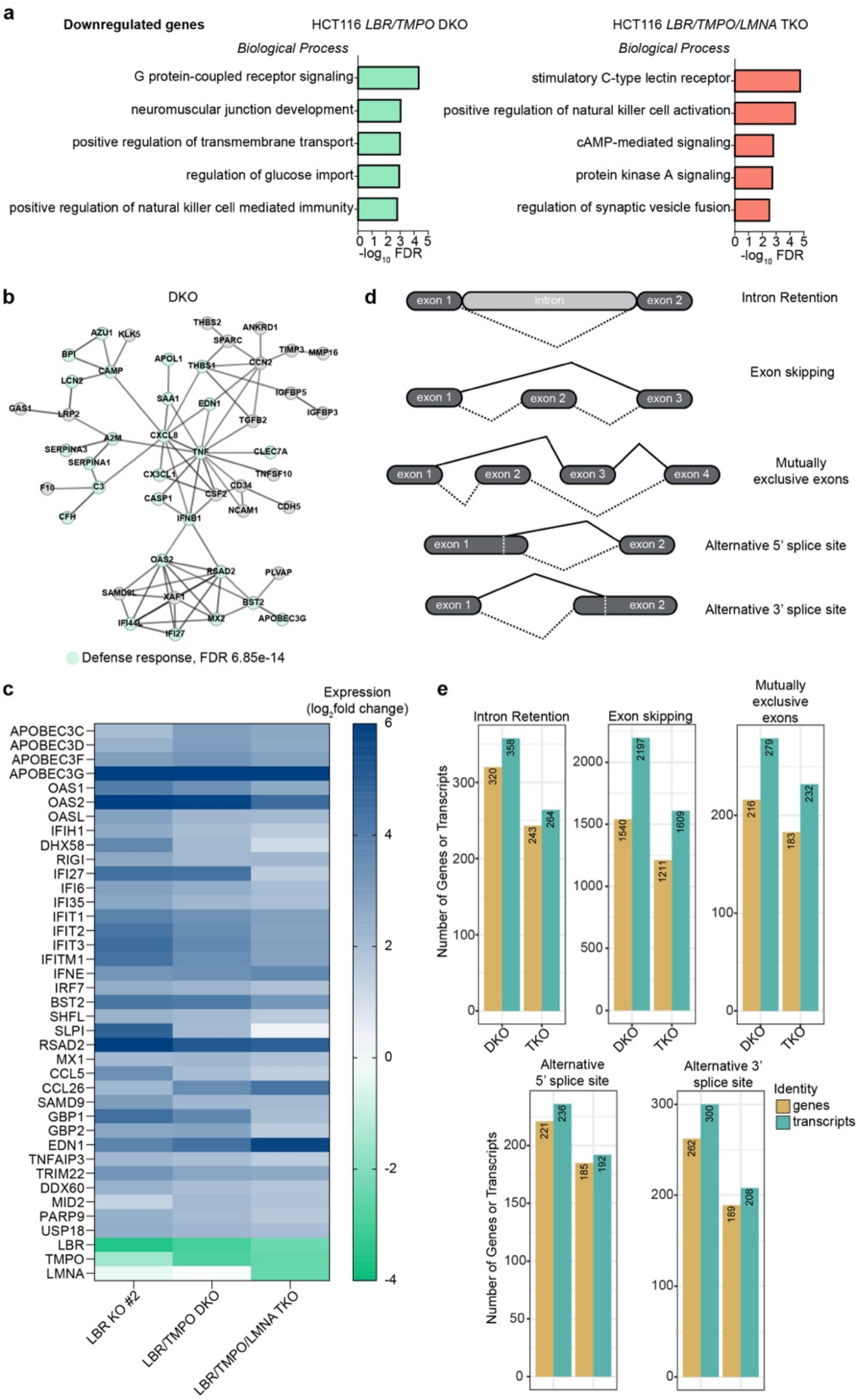


**Figure S7. The loss of heterochromatin tethers leads to an upregulation of RNA innate immunity genes and splicing defects in HCT116 cells.**

**a**, Bar graph of GO term analysis ('Biological process' category) of significantly downregulated genes with a $\log_2$-fold change ≤ -2 in HCT116 *LBR/TMPO* DKO and *LBR/TMPO/LMNA* TKO cells. Fischer's exact test was used to determine overrepresented terms.

**b**, K-means clustering of the top 200 upregulated protein coding genes in the *LBR/TMPO* DKO cells into a functional cluster. The genes within the cluster relating to the 'defense response' GO term were highlighted in green. Connections between genes correspond to high confidence interactions based on the STRING database ($p > 0.7$)[129].

**c**, Heatmap of the mean expression of upregulated RNA innate immunity genes identified by GO terms 'negative regulation of viral genome' and 'defense response to virus' for the indicated KO backgrounds.

**d**, Schematic depiction of the analysed differential splicing defects.

**e**, Bar plots depicting number of genes and transcripts identified in different deregulated splicing events. Events with IncLevelDifference > |0.15| and FDR < 0.05 were considered significant.

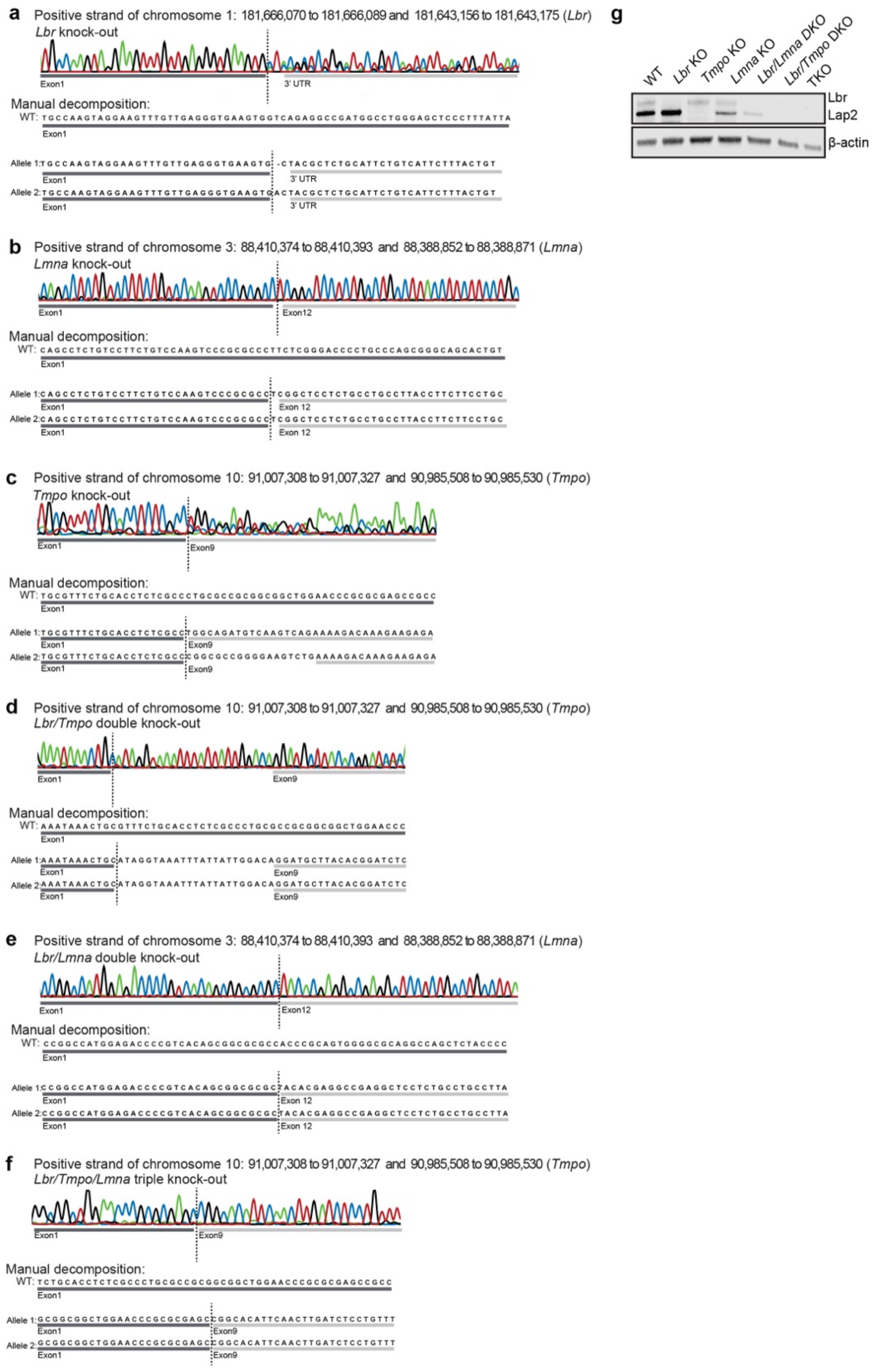


**Figure S8. Characterisation of mESCs knock-out cell lines.**

**a**, Characterisation of the E14TG2a *Lbr* KO. Genomic coordinates correspond to the locations that guides 1 and 2 bind. Manual decomposition of the sequence of the *Lbr* gene confirms deletion of the gene in both alleles.
**b** Characterisation of the E14TG2a *Lmna* KO. Manual decomposition of the sequence of the *Lmna* gene confirms deletion of the gene in both alleles.
**c**, Characterisation of the E14TG2a *Tmpo* KO. Manual decomposition of the sequence of the *Tmpo* gene confirms deletion of the gene in both alleles.
**d**, Characterisation of the E14TG2a *Lbr/Tmpo* DKO. Manual decomposition of the sequence of the *Tmpo* gene confirms deletion of the gene in both alleles. This KO was performed on the *Lbr* KO background.
**e**, Characterisation of the E14TG2a *Lbr/Lmna* DKO. Manual decomposition of the sequence of the *Lmna* gene confirms deletion of the gene in both alleles. This KO was performed on the *Lbr* KO background.
**f**, Characterisation of the E14TG2a *Lbr/Tmpo/Lmna* TKO. Manual decomposition of the sequence of the *Tmpo* gene confirms deletion of the gene in both alleles. This KO was performed on the *Lbr/Lmna* DKO background.
**g**, Immunoblot of Lbr and Lap2 in the various knock-out backgrounds to validate the loss of these proteins.

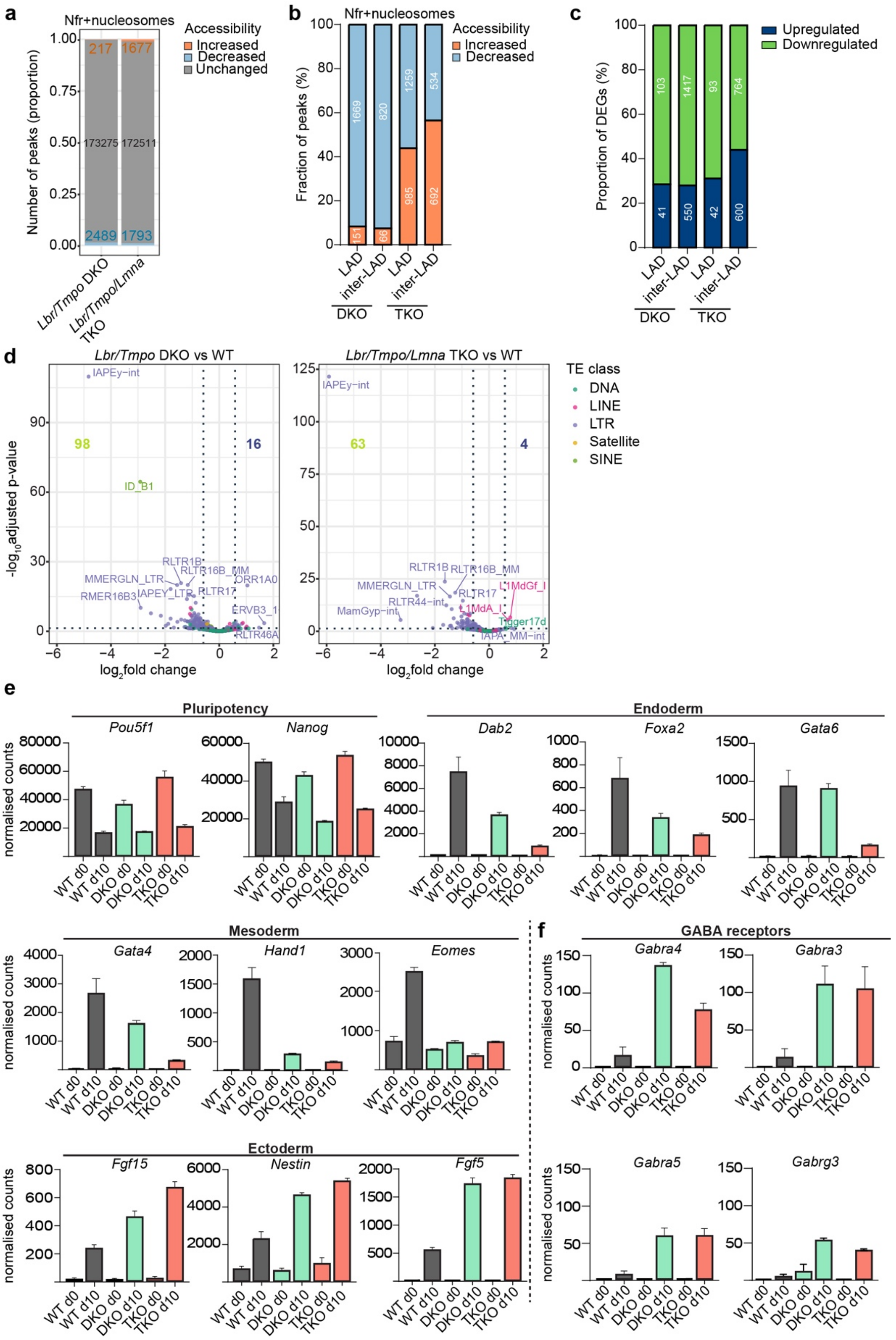


**Figure S9. Changes in the transcriptome, chromatin organisation and cell differentiation upon the loss of heterochromatin tethers in mESCs.**

**a**, Proportion of ATAC-seq peaks in Nfr+nucleosomes regions with increased or decreased accessibility in *Lbr/Tmpo* DKO and *Lbr/Tmpo/Lmna* TKO mESCs (N = 3). All loci with a $log_2$-fold change ≥ I1I and an adjusted p-value ≤ 0.05 were considered significant.

**b**, Bar graph highlighting the proportion of Nfr+nucleosomes ATAC-seq peaks, with increased or decreased accessibility in regions originally designated as LADs in mESCs[74], for both *Lbr/Tmpo* DKO and *Lbr/Tmpo/Lmna* TKO mESCs.

**c**, Bar graph showing the fate of deregulated genes (identified by RNA-seq) previously positioned in LADs and inter-LADs in *Lbr/Tmpo* DKO and *Lbr/Tmpo/Lmna* TKO mESCs.
**d**, Volcano plots of transposable elements in E14TG2a *Lbr/Tmpo* DKO and *Lbr/Tmpo/Lmna* TKO vs. E14TG2a WT cells (N = 3). Genes with an adjusted p-value ≤ 0.05 are marked as up- or downregulated if log2-fold change ≥ I I0.585I.
**e**, Bar graphs of normalised read counts in RNA-seq of marker genes for pluripotency and the three germ layers in E14TG2a WT, *Lbr/Tmpo* DKO and *Lbr/Tmpo/Lmna* TKO cells after 10 days of embryoid body differentiation (N = 3). Error bars, SD.
**f**, Bar graphs of normalised read counts in RNA-seq of select GABA receptor genes in E14TG2a WT, *Lbr/Tmpo* DKO and *Lbr/Tmpo/Lmna* TKO cells after 10 days of embryoid body differentiation (N = 3). Error bars, SD.

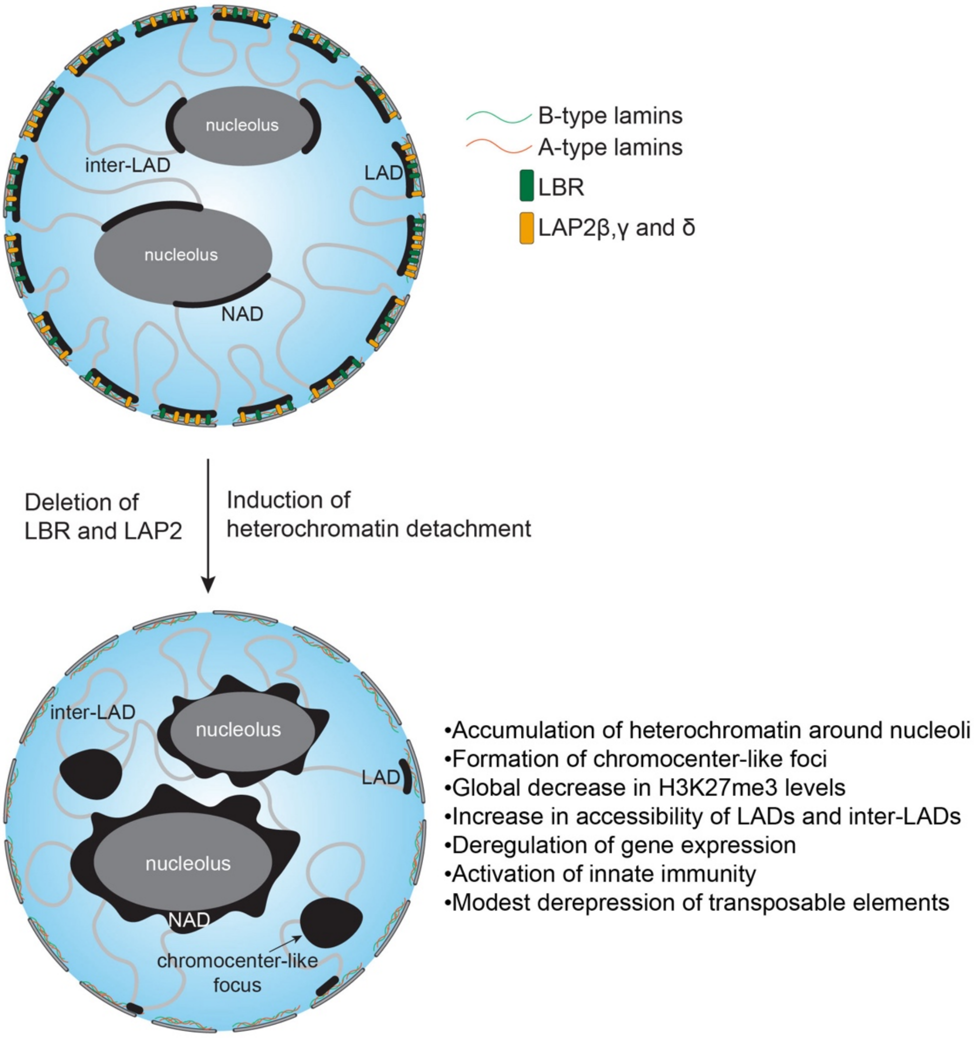


**Figure S10. Model of nuclear reorganization upon deletion of NE tethers in differentiated and pluripotent mammalian cells.**

The loss of LBR and LAP2 causes a global reduction of heterochromatin at the NE, accompanied by repositioning of heterochromatin into the vicinity of nucleoli and the formation of chromocenter-like foci within the nuclear interior. In *LBR/TMPO* KO cells, loss of tethering is ultimately associated with major changes in chromatin accessibility throughout the genome, a reduction in the global levels of H3K27me3, deregulation of the transcriptome, the activation of innate immunity genes, a modest activation of transposable elements (observed only in HCT116 cells) and defects in stem cell differentiation.
